# Social cues on stone tools outweigh raw material properties in wild primates

**DOI:** 10.1101/2024.06.10.598233

**Authors:** J Henke-von der Malsburg, J Reeves, T Proffitt, T Falótico, HP Rufo, LV Luncz

## Abstract

The ability to select appropriate tool material enabled early hominins access to new resources and environments. The underlying mechanisms driving tool selection effectively remain unknown. Observations of extant primates have demonstrated strong selectivity for specific tools, offering analogous insight into technological decision-making. However, whether tool selection is determined by individual experience alone or social information plays a role remained difficult to disentangle. Here, we used an experimental approach to investigate decision-making factors for tool selection in non-human primates. We provided naturalistic nut-cracking opportunities to wild capuchin monkeys, one of the most prolific extant tool users. We offered standardized stones varying in asocial (material properties) and social cues (evidence of previous use) to two populations, differing in their previous experience of natural materials. Our results show that both populations persistently selected tools based on their material properties when only asocial cues were provided. However, when provided with both asocial and social cues combined, they consistently selected previously used material regardless of material properties. These findings suggest that wild capuchin monkeys discriminate between raw material properties; however, prioritize social cues when present. Tool selection behaviors are therefore shaped by indirect social processes and highlight the importance of culturally transmitted information for skill acquisition in technological primates.

## 2. Introduction

The use of tools has shaped the course of human evolution, allowing access to a wider range of environments and resources. A fundamental requirement of tool use may be the capacity to select materials suitable for the task. Selection behavior is thought to require an interplay of cognitive, ecological, and cultural processes^1,2^. Proficient tool manufacturing and use requires the cognitive capacity to recognize and select materials with physical properties that are task-appropriate^3–5^. Ecological factors impose constraints on tool use by influencing the spatial distribution and abundance of materials suitable for tool-using tasks^6,7^. Culture contributes to shaping the acquisition and maintenance of tool use through the social transmission of knowledge surrounding selection criteria^8–10^. Although there is a general understanding of how ecology and cognition influence selection, little is known about how social processes influence such behaviors and, more broadly, the evolution of tool use.

As past behavior of hominins can only be inferred through recovered stone artefacts, extant stone-tool-using primates can serve as valuable analogs for understanding the interplay of factors driving tool selection^11,12^. Such opportunities allow us to observe tool use within its broader socio-ecological context^13^. Here direct observations allow us to establish connections between social behavior and environmental factors that influence tool selection and use. While living primates cannot serve as stand-ins for our hominin ancestors, understanding interactions between their tool use behaviors, and broader socio-ecological factors allow us to generate robust hypotheses about the potential mechanisms behind the evolution of tool-use.

Studies of Western chimpanzees (*Pan troglodytes verus*)^14^, Burmese long-tailed macaques (*Macaca fascicularis aurea*)^15^ and capuchin monkeys (*Sapajus* spp., *Cebus capucinus imitator*)^16–19^ show that these species select tool materials based on physical properties, such as size, weight, and shape, best suited for the task^20–24^. For example, the anvil and hammerstone properties used to process certain nut species covary with the hardness, shape, or chemical characteristic of the nuts^14,25–28^. At Fazenda Boa Vista, where capuchin monkeys (*Sapajus libidinosus*) crack relatively hard and large palm nuts (*Astrocaryum campestre*, *Attalea barreirensis*, *Orbignya* sp., and *Attalea* sp.^29^), individuals preferentially select heavier hammerstones used in conjunction with anvils used in prior nut-cracking activities. These anvils exhibit pits from previous nut-cracking, contributing to the stabilization of the round palm nut^30,31^. In addition, ecological studies have demonstrated that the frequency of stone tool use is heavily dictated by the availability of suitable stone materials combined with their proximity to food sources^6,32^.

With respect to culture, some studies suggest that tool selection is shaped through a combination of social learning mechanisms, including observational learning and local enhancement^8,33,34^. The selection, transport, and discard of tools can relocate accumulations of previously used materials away from the source, modifying the accessibility of tool materials and the number of tool-use sites available^27,32^. These accumulations of discarded tools not only serve as sources of stone for future tool use events^1^ but may serve as additional learning opportunities, providing novice individuals with opportunities to interact with materials in the context in which they are used as tools^35^. These interactions may influence learning and stone selection. The traces of previous use found on discarded tools or patterns of discarded food may act as valuable socially mediated cues, facilitating visual associations between specific tasks and raw materials and, consequently conditioning the selection choices of novices. In capuchin monkeys, observational studies report younger individuals observing older conspecifics cracking nuts and re-using previous tools, creating specific social learning opportunities^36–38^.

Here, we investigate the social factors of tool selection through a systematic study of two populations of bearded capuchin monkeys (*Sapajus libidinosus*) using an experimental approach in their natural environment. Both populations crack nuts using anvils and hammerstones as a part of their daily foraging behaviors (Figure 1A)^39,40^. To investigate the influence of social factors on stone selection during nut-cracking we provided subjects with three nut-cracking tasks. These tasks varied in the material properties of the stones provided, the visible signs of previous use for nut-cracking (percussive damage on stones, and the presence of nut residues), or the arrangement of the stones. In the first task (Task 1), we tested whether individuals were able to select anvils and hammerstones according to their material property (hardness). After we established that capuchin monkeys differentiate between raw material, in Tasks 2 and 3, we tested whether social cues (signs of previous use) influence the choice of foraging site and tool selection. Across all tasks, we investigated, how selection preferences influenced foraging efficiency over time, as well as their consistency across study groups.

**Figure 1.**
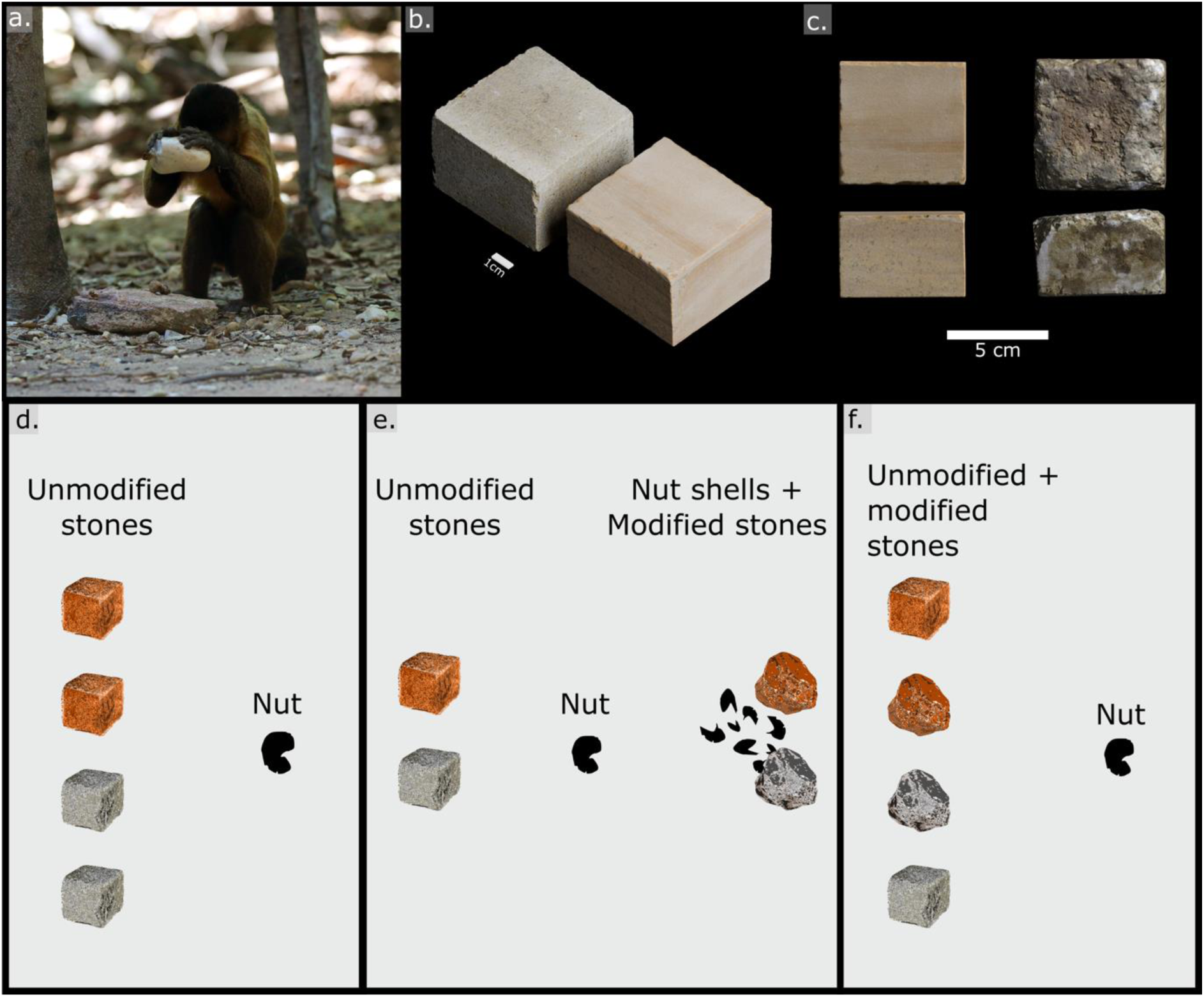
a. A capuchin monkey cracking a cashew nut using natural stones. b. Visual comparison of the colors associated wirh each materials provided at nut-cracking sites, c) Examples ofmodified sandstone and modified limestone. d.-f. Experimental tasks (c. Task 1, d. Task 2, e. Task 3) offering nut-cracking sites with stones of different materials (depicted by different colors) and modification (depicted by shapes)

## 3. Results

To assess whether capuchin monkeys use asocial or social attributes for their selection of anvils and hammerstones, we employed a three-task experimental design (Figure 1C-E). In each task, we provided naturalistic foraging sites with standardized stones of two raw materials unknown to the capuchin monkey groups and locally available nuts (cashew nuts at Serra da Capivara National Park (SCNP) and palm nuts at Parque Ecológico do Tietê (PET)). Whenever we encountered the capuchin monkeys, we cleared the area of any other stone material to ensure that only provided stones were used. In Task 1, we tested the ability to distinguish between materials of different hardness. We provided two unmodified limestone and sandstone blocks of a standardised shape and mass in a randomized order. In Task 2, we provided two potential nut-cracking sites, one of which was modified to display clear signs of previous use (social cues), including the presence of broken nut shells and percussed stones (one limestone and one sandstone). The other contained only one unmodified, standardized limestone and sandstone block (as in Task 1). Task 3 tested the influence of stone modification (percussed vs. unmodified stones) on material selection by providing two limestone and two sandstone blocks, of which, one of each showed visual signs of previous use (damaged surfaces and nut residue), with the other two remaining unmodified. Individuals interacted with the setups voluntarily during their daily foraging routes. Subjects were exposed to the tasks in successive order, starting with the following task only when 5‒10 trials had been completed by each participating individual. A total of 414 trials were conducted over the three experimental tasks which included the participation of 12 wild and 15 semi-wild capuchin monkeys from SCNP and PET, respectively (SI Table S1).

The outcomes of these experiments are reported as a series of generalized linear mixed models (significance of full-null model comparisons, effect sizes *R²*) and the influence of the stone material, their percussive modification (Figure 1B), the study group, and the nut-cracking performance (significance of full-reduced model comparisons, estimates *E*) are reported. In each reported model, we controlled for the subject’s sex, the continuous experimental participation, as well as random effects of stone material, stone ID, trial ID, and focal subject ID (see SI methods). A detailed description of all experimental designs, the testing procedure used, and the statistical models employed (full-model descriptions, significance testing, model diagnostics) along with the full results are provided in the supplementary online material.

### 3.1. Capuchin monkeys discriminate between stones based on material properties

When provided with a choice of two standardized blocks of harder sandstone, and two blocks of softer limestone as nut-cracking tools (N = 152 trials; Task 1, Figure 1C), the material had a significant effect on the subjects’ selection for both anvils, (model 1a: *X²*(4) = 15.03, *p* = .022; *R²* = .04; SI Table S2) and hammerstones (model 1b: *X²*(4) = 1.56, *p* = .004; *R²* = .02; SI Table S3). Capuchin monkeys selected the softer limestone as anvils more frequently than the harder sandstone (model 1a: *E*_sandstone_ = ‒0.64 ± 0.24, *p* = .002; Figure 2A; SI Table S4). Conversely, the harder sandstone was more often selected as hammerstones compared to the softer limestone (model 1b: *E*_sandstone_ = 0.82 ± 0.32, *p* = .001; Figure 2A; SI Table S4). This suggests that the hardness of raw material plays a substantial role in the selection patterns of these tool types for capuchin monkeys.

**Figure 2.**
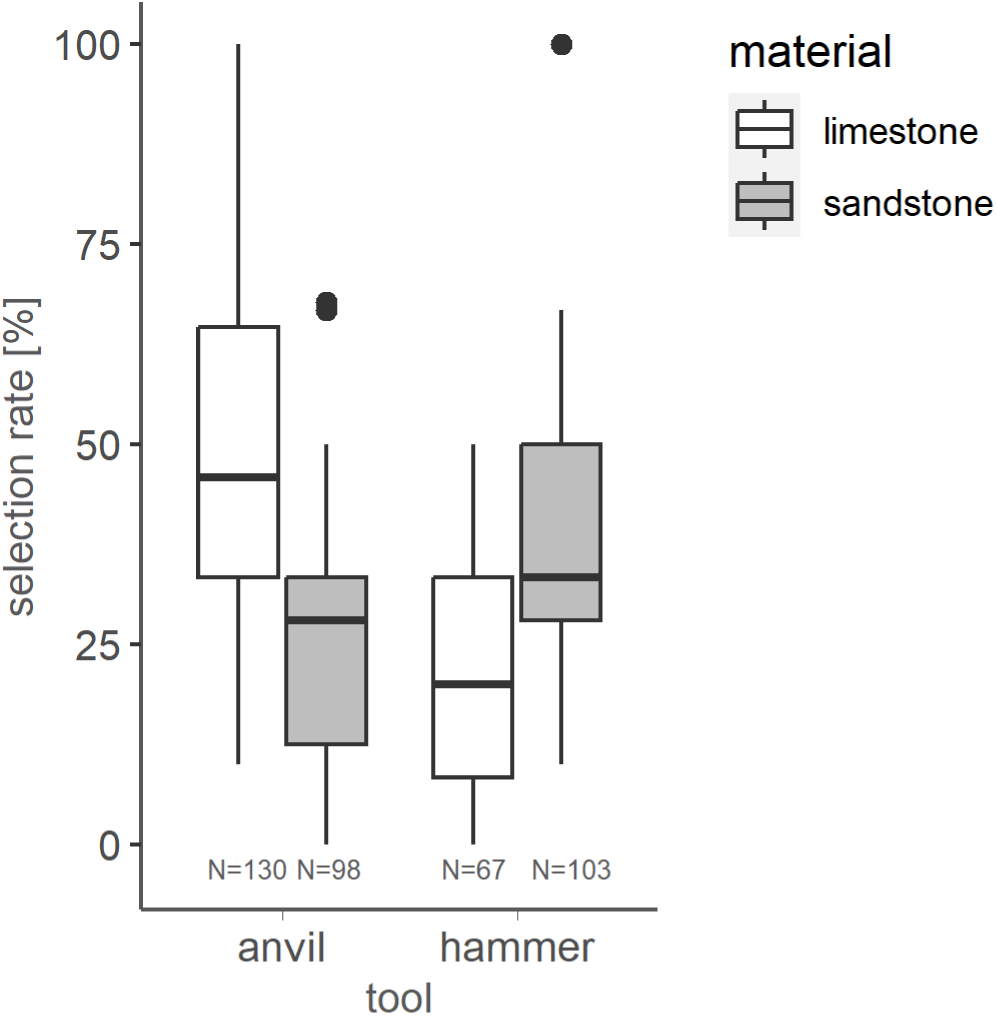
Task 1: Selection rate of raw materials for anvils and hammerstones. Capuchin monkeys rather selected softer limestone anvils (binomial linear mixed model, *p* = .002) and harder sandstone hammers (binomial linear mixed model, *p* = .001). Shown are median and standard error bars. Sample sizes are given below each boxplot

### 3.2. Capuchin monkeys select previously used nut-cracking sites

When provided with a choice of two separate nut-cracking sites (N = 221 trials; Task 2, Figure 1D),the presence of previous nut-cracking activities had a significant effect on site selection (model 2a: *X²*(4) = 14.19, *p* = .007; *R²* = .05; SI Table S5). Capuchin monkeys preferentially chose nut-cracking sites that bear traces of previous use compared to sites containing only unmodified stones (*E*_unmodified_ = ‒1.86 ± 0.40, *p* <.001; Figure 3A), suggesting that local enhancement plays a crucial role in initial nut-cracking site selection.

**Figure 3.**
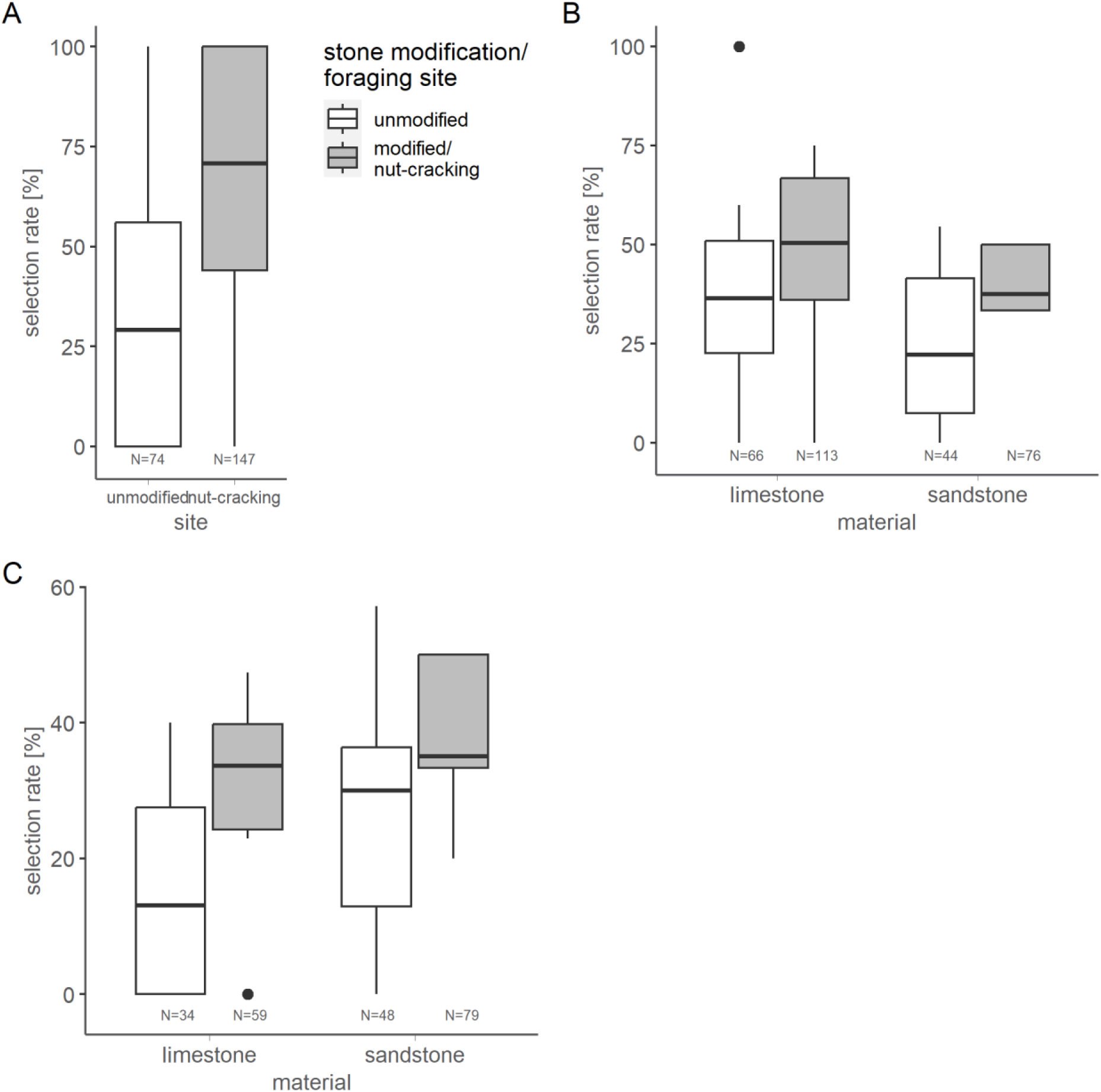
Task 2: Selection rates for foraging sites (A), anvils (B), and hammerstones (C). Subjects first oriented towards previously used nut-cracking sites (binomial linear mixed model, *p* <.001). Then, they rather selected modified stones (grey), softer limestone anvils (binomial linear mixed model, *p* = .001) but harder sandstone hammers (binomial linear mixed model, *p* = .022). Shown are median and standard error bars. Sample sizes are given below each boxplot

Furthermore, capuchin monkeys showed a clear preference to also use modified stones as both anvils (model 2b: *X²*(5) = 23.87, *p* = .001; *R²* = .08; N = 189 trials; SI Table S6) and hammerstones (model 2c: *X²*(5) = 6.81, *p* = .016; *R²* = .03; N = 189 trials; SI Table S7) whilst maintaining the same preference for raw material properties. They continued to select softer limestone to use as anvils (model 2b: *E*_sandstone_ = ‒0.91 ± 0.20, *p* = .001; Figure 3B; SI Table S4) and harder sandstone as hammerstone (model 2c: *E*_sandstone_ = 0.68 ± 0.34, *p* = .022; Figure 3B; SI Table S4).

### 3.3. Capuchin monkeys discriminate between stone tools based on social cues rather than material properties

Finally, in task 3 (N=73, Figure 1E), when one sandstone block was modified to mimic traces of previous nut-cracking activities, the subjects repeatedly discriminated between these stones for their anvil (model 3a: *X²*(5) = 5.00, *p* = .004; *R²* = .07; SI Table S8) but not hammerstone selection (model 3b: *X²*(5) = 2.63, *p* = .107; *R²* = .02; SI Table S9). Unlike the previous tasks, no single raw material was preferentially selected based on their properties (Anvils: model 3a: *E*_sandstone_ = ‒0.37 ± 0.35, *p* = .255; Hammerstones: model 3b: *E*_sandstone_ = 0.50 ± 0.37, *p* = .247; Figure 4A; SI Table S4). Instead, modified stones were more frequently selected as anvils (model 3a: *E*_modified_ = 1.08 ± 0.30, *p* = .003; Figure 4B). However, no preferential selection was observed for hammerstones, (model 3b: *E*_modified_ = ‒0.65 ± 0.39, *p* = .112; Figure 4B).

**Figure 4.**
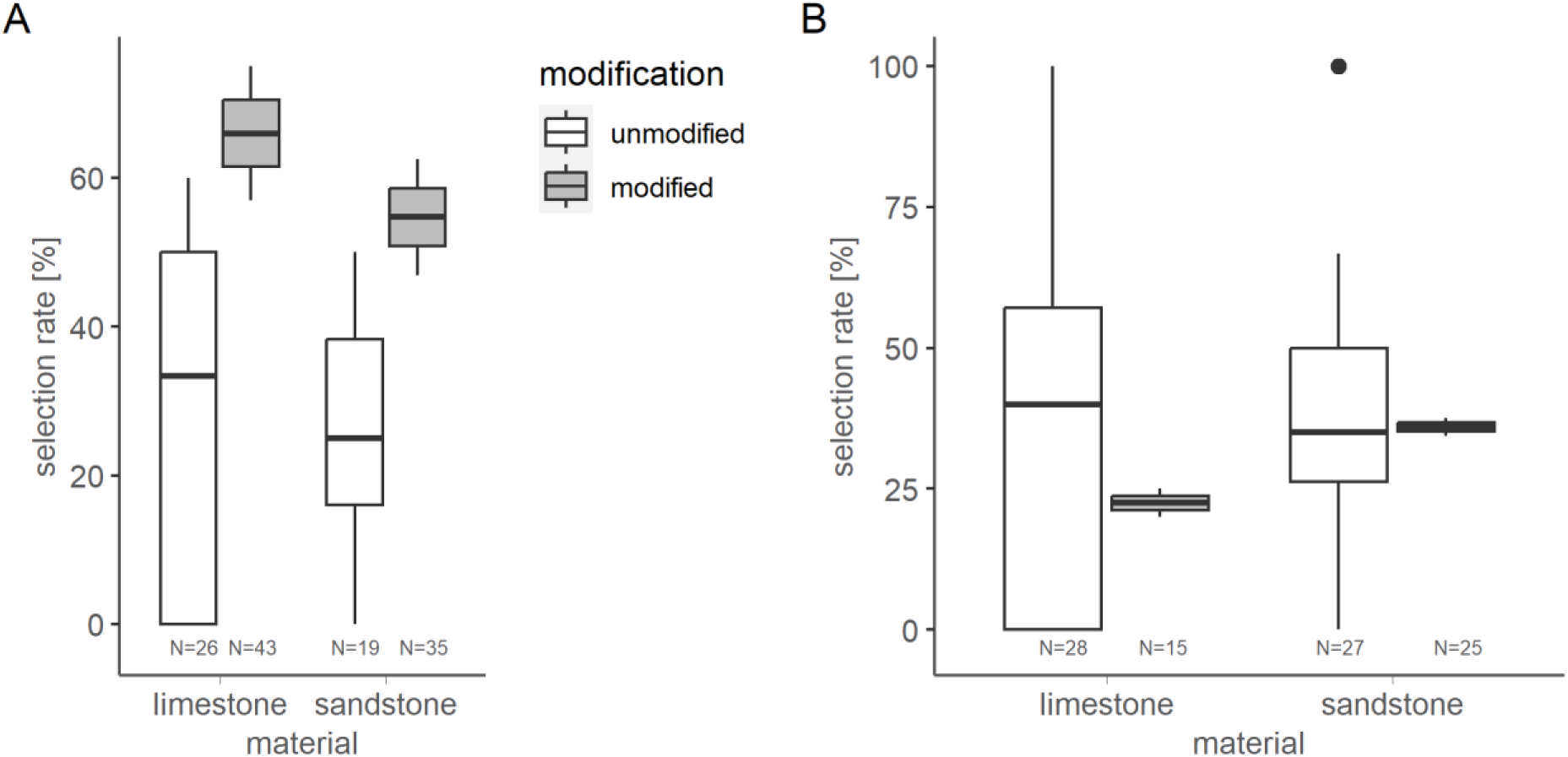
Task 3: Selection rates for anvils (A) and hammerstones (B). As anvils, subjects rather selected modified stones (grey; binomial linear mixed model, *p* = .003), irrespective of the material (*p* = .255). As hammers, subjects did not discriminate between materials or stone modifications (binomial linear mixed model, *p* = .107). Shown are median and standard error bars. Sample sizes are given below each boxplot

### 3.4. Stone tool selection patterns did not affect the nut-cracking performance across capuchin populations

Despite the clear preference for specific raw material properties for both anvils and hammerstones, this selection pattern had little effect on foraging efficiency. Instead, the probability of success was influenced more by the subjects’ individual characteristics (model 4a: *X²*(5) = 22.28, *p* <.001; *R²* = .08; N = 405 trials; SI Table S10), with capuchin monkeys from SCNP being more likely to succeed than those from PET (model 4a: *E*_SCNP_ = 1.54 ± 0.43, *p* <.001; Figure 5A). The primary cause of failure for PET individuals was loss of the nut during nut-cracking leading to the subject abandoning the experimental site. At SCNP, individuals primarily left the experimental site with the nut, continuing nut processing at natural anvils (Table S11). In both groups, individual success probability increased across trials (*E* = 0.51 ± 0.17, *p* = .005; **Error! Reference source not found.**B), suggesting the presence of a learning curve for handling previously unknown raw material. However, the materials of the chosen stone tools did not affect the success probability (anvil: *E*_sandstone_ = 0.04 ± 0.40, *p* = .920; hammerstone: *E*_sandstone_ = 0.16 ± 0.34, *p* = .632). The nut-cracking efficiency in successful trials was not influenced by any of the subjects’ or stone characteristics (Nut-cracking speed, model 4b: *X²*(5) = 2.15, *p* = .826; *R²* = .34; N = 292 trials; SI Table S12; Number of strikes, model 4c: *X²*(5) = 4.18, *p* = .525; *R²* = .01; N = 292 trials; SI Table S13).

**Figure 5.**
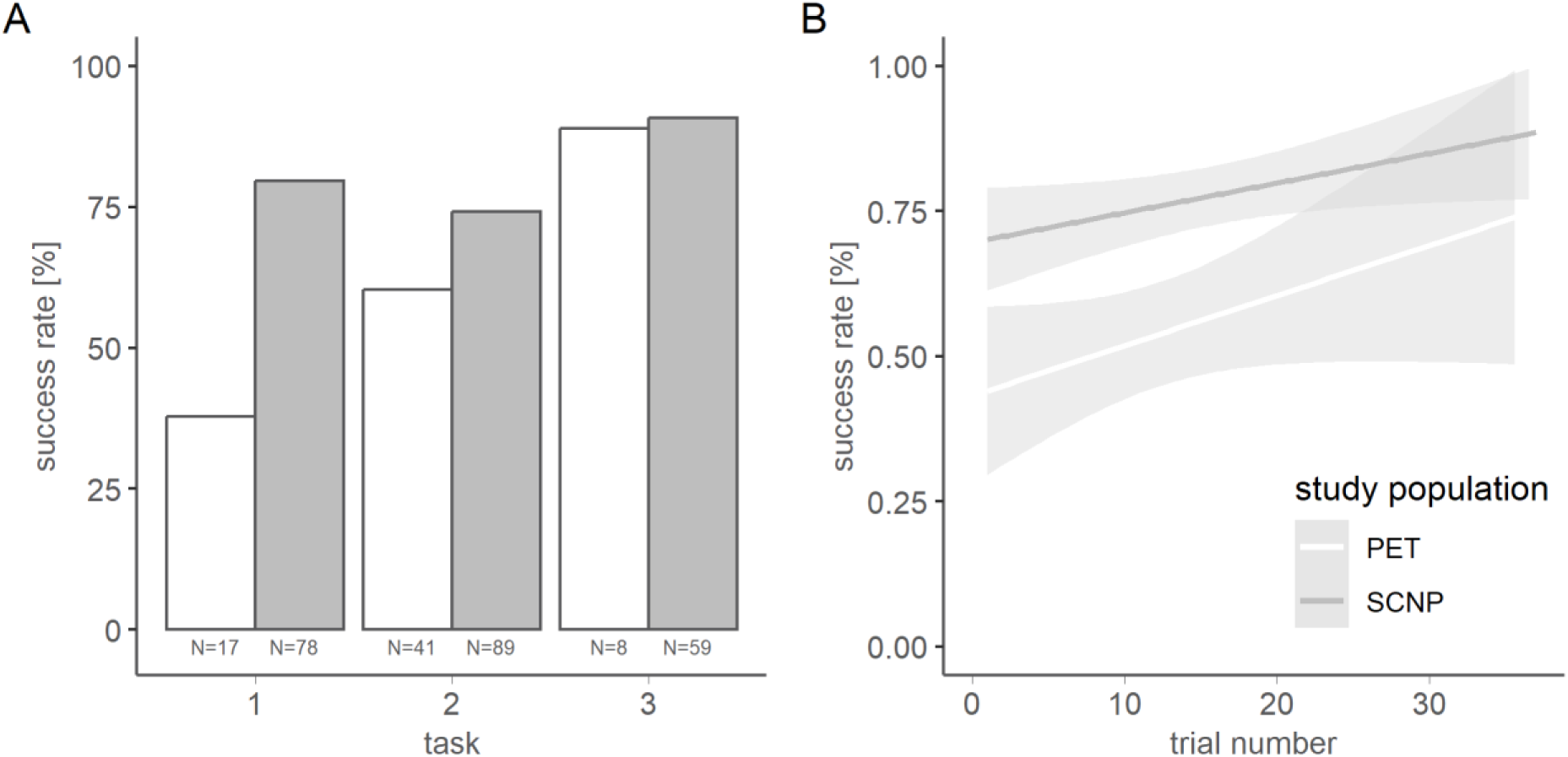
A Success rates in the individual tasks and B success probabilities across trials for capuchin monkeys at SCNP (grey) and PET (white). Individuals at SCNP were more successful than individuals at PET (binomial linear mixed model; *p* <.001). Individuals of both populations were more successful in later trials (*p* = .005). Sample sizes are given below bar plots

## 4. Discussion

Studying the material selection associated with tool use provides insight into the cognitive abilities of tool users. Both early hominins and nonhuman primates have shown evidence of systematic exploitation of raw materials, leading to landscape-wide modifications of material assemblages^1,41–47^. However, how socially mediated information embedded at previously used tool sites influences selection is less clear. Offering naturalistic nut-cracking sites with standardized stones varying in physical properties of stone, and social factors, such as evidence of previous use, this study sought to investigate decision-making for anvil and hammerstone material in capuchin monkeys.

Our findings highlight a complex interplay between asocial and social factors related to tool selection and provide insight into the cognitive capacities of nut-cracking primates. In the absence of social cues, capuchin monkeys demonstrate the ability to select nut-cracking tools by distinguishing between raw materials that vary in hardness. Specifically, softer stones are consistently preferred as anvils, while harder stones are selected as hammerstones. This observation implies that capuchin monkeys differentiate between raw materials based on their hardness. While previous studies have shown that primates select percussive tools according to both their geometric attributes^27^, and the attributes of the target object^28^, this result indicates that capuchin monkeys also are discerning of material properties in their selection process.

Previous studies have shown that raw material type can affect the efficacy of percussive stone tools^28,48^. Specifically, the use of a softer raw material has been found to promote the gradual development of a pit on the surface of the anvil, arising due to nut cracking. By increasing the stability of the target nut, this wear pattern could enhance the efficiency of this behavior ^49^. However, this enhancement is only pronounced with round nuts (as seen with palm nuts provided at PET) but may not be as beneficial for flat, elongated cashew nuts (provided at SCNP). As a result, palm nuts at PET were more prone to dislodging from the anvil, leading to individuals to cease further nut-cracking (Table S11). Despite this, the potential advantage of a stabilizing effect may suggest a more robust selection effect for previously used anvils compared to hammerstones. Conversely, the use of soft hammerstones, which absorb force or deform through use, is likely to be less efficient compared to harder materials that transmit a larger portion of the tool users force onto the nut. This higher rate of force transmission, theoretically, could reduce the number of strikes required to successfully crack the nut.

Given that the stones used in our study were standardized in terms of shape and dimensions, how capuchin monkeys differentiate between hardness remains unclear. Although the weight difference between the stones was minor (SI Table S14), the monkeys may have relied on weight as a determining factor in their choice, given that the harder tools were slightly heavier, and capuchin monkeys have demonstrated their ability to assess weight^50,51^. However, they never lifted stones without subsequently using them as anvils or hammerstones, indicating that it was unlikely that they evaluated stone weight prior to use. Our observations suggest that capuchin monkeys may have been frequently assessing the material properties through more subtle exploratory behaviors, such as tapping (see supplementary Video 1), a behavior that capuchin monkeys also use to assess the ripeness of uncracked nuts^50^.

However, when two potential nut-cracking sites were offered (Task 2), both of which were equipped with suitable stone materials, the monkeys consistently selected the one with clear evidence of previous successful nut-cracking activity. This result suggests that socially mediated factors, such as the residual traces of previous activities, influence the stone selection process. The presence of both visible nut debris and used tools together may encourage the reuse of these sites compared to unused localities. Consequently, the repeated use of these sites, coupled with their increasing visibility, potentially reinforces ongoing tool-related behaviors. This, in turn, may contribute to the persistence of nut-cracking practices within primate populations once established. This result supported findings of previous studies^35^ that social cues of tool use embedded in the environment contribute to shaping knowledge about the selection of tool materials.

The importance of social cues in shaping tool selection is further supported by the results of our third experiment. In this experiment tools could be selected from a sample of unmodified and modified stones. Here, capuchin monkeys based their choice of anvil on the prior modification of the stone rather than the raw material itself. This preference for selecting materials modified by previous behaviors supports the hypothesis that the social context of tool use acquisition may extend beyond direct observation.

Our findings further allow us to discuss the implications of tool selection for cultural evolution. One of the main tenets of cultural evolutionary theoy is that individuals do not only inherit their genes but also the environments and social practices of their ancestors^52^. In this light, the combination of discarded tools and residual nut debris may serve as learning opportunities for future generations. Therefore, tool accumulations may help to reinforce the selective preferences of the older more skilled nut-crackers in those learning, creating path-dependent behaviors within a given group. Such a process would not only reinforce the types and attributes of materials individuals associate with suitable tools over time but also may result in group-specific behaviors across species^35–37,53^.

Our results contribute to a growing body of evidence that tool use is guided by social cues. Previous research has shown that female chimpanzees (*Pan troglodytes versus*), upon immigrating into a new group, exhibit a propensity to conform to the tool selection patterns practiced by the new host group members^54,55^. As such, their previous tool use preferences are relinquished. It is thought that conformity to local traits allows primates to quickly adopt the most efficient behavior within a specific environmental context^56,57^, enhancing their foraging success, overall fitness, and survival.In addition to chimpanzees, different social groups of wild long-tailed macaques (*Macaca fascicularis*) show distinct selection patterns concerning stone tool properties^53^. This group’s coherent selection patterns, particularly in the form of visually identifiable percussive damage on stone tools, highlight the potential importance of social cues across the range of tool-using primate species.

Furthermore, our results concerning stone selection decisions were persistent across study groups, despite different experiences with natural materials (stones as well as nuts). However, nut-cracking proficiency varied between the different populations of capuchin monkeys, with wild individuals at SCNP exhibiting higher success rates in nut-cracking compared to the semi-wild individuals from PET. The respective nut-cracking skills are likely shaped by a combination of individual, environmental, and social factors, as well as differences in mechanical properties of the presented nuts^28^. Inconsistent significant effects of the study population, sex and trial (i.e., specific experimental experience) on stone selection decisions and nut-cracking performance further suggest that individual factors (e.g., population-specific motivation) contribute to variances in nut-cracking activities. For example, individuals at PET were found to switch more often between stones within one trial, whereas individuals at SCNP usually continued to crack nuts with the selected anvil and / or hammer once they chose a specific stone. This may imply variation in population-specific motivation for nut-cracking, resulting from wild individuals at SCNP being challenged by reduced food availability during the dry season, while semi-wild individuals at PET were provisioned with additional high-quality food (e.g., fruits). This may explain why, SCNP individuals were more efficient nut-crackers, than PET individuals. Additional effects of sex may reflect variances in individual experience either in nut-cracking or within the experiments, as well as variances in social learning opportunities, which has been investigated in more detail in previous studies^58,59^. For instance, male capuchin monkeys were found to use tools more frequently than females, suggesting more experience, although nut-cracking efficiency did not vary between sexes^58^.

Despite the evident preference displayed by capuchin monkeys for specific tool materials (asocial cues) and indicators of prior use (social cues), we found no significant effects of tool selection on the success or efficiency of nut-cracking. Since the participating individuals were already proficient nut-crackers, their extensive experience likely overshadows improvements in nut-cracking efficiency due to material selection. Since all material combinations could lead to nut-cracking success, a learning effect that a certain combination (e.g. soft anvil, hard hammer) was more efficient than another (e.g. hard anvil, soft hammer) may be hidden. Alternatively, the monkeys learned over the course of the experiments that they would succeed no matter which material combination they chose, resulting in the elimination of initial preference (Task 1 compared to Task 3). Instead, the monkeys rather based their stone selection on the presence of social cues indicating prior use which we interpret as more important factor for experienced nut-crackers than the mere stone material in our experiment.

Beyond our understanding of primate tool selection, the results of this study also have implications for interpreting hominin selection, where the social context of tool production is often debated. For example, the repeated use of specific locations in the landscape is also a characteristic feature of the Oldowan technological complex observed in Plio-Pleistocene hominins^43,60,61^ as well as more recent forager communities^62^. Previous research has emphasized the role of landscape modification through repeated tool activities as a potential contributing factor to the recurrent use of favored locations by Plio-Pleistocene hominins^43,63^. Once a behavioral location became visually distinguishable due to the material presence or modification by hominins, it may, itself have served as a powerful visual and social cue to both contemporaneous and future hominin groups, highlighting the suitability of the location and raw material^64^. Given the limited availability of direct behavioral and cognitive evidence associated with past hominin species, there is little data to directly support this hypothesis. Nonetheless, our results indicate that the visual identification of successful past behaviors and respective discrimination between raw materials can motivate the reuse of specific locations and materials in the landscapes.

The ability to identify and exploit favorable material properties for specific tasks is also a fundamental characteristic of early hominin tool behaviors. During the Plio-Pleistocene, Oldowan hominins routinely displayed a preference for high-quality isotropic stones with advantageous natural morphologies for stone tool production^46,65,66^. Our study suggests that the ability to discern stone material properties may be a shared capacity across tool-using primates, including hominins, regardless of phylogenetic divergence and drives primate technological culture.

## 5. Materials and Methods

### 5.1 Study animals and study sites

This research was conducted with two populations of bearded capuchin monkeys (*Sapajus libidinosus*), a wild group from Serra da Capivara National Park (SCNP), in Piauí, and a semi-wild group in the Park Ecológico do Tietê (PET), São Paulo, both which are in Brazil. Both populations are habituated to observational and experimental studies^27,67–70^.

Serra da Capivara National Park is a semi-arid region characterized by dry scrub vegetation and deciduous forest^71^. Available stones in this area predominately include natural quartzite cobbles (recrystallized quartz and polycrystalline quartz with metamorphic foliation and high silica content) and tabular sandstones blocks, as well as concrete associated with artificially built pathways^72,73^.

Capuchins monkeys (*S. libidinosus* and *S. nigritus*) at the PET, are free to roam both, within and outside the park but are food-provisioned at a visitor-restricted area by the Centro de Recuperação de Animais Silvestres (CRAS) staff. Within this area, nut-cracking sites are located around palm trees (*Syagrus romanzoffiana*). The available stone tool material at PET differs from that found at SCNP as it is mostly associated with the anthropogenic history and modification of the park. As such, the most commonly raw material used for nut-cracking hammerstones includes concrete and granite blocks, as well as natural sandstone and ironstone, while concrete pavements are predominately used as anvils.

Given these differences in naturally available material (stones and nuts), the populations entered the study differing in their nut-cracking experience, as well as motivational aspects arising from seasonal food availability (SCNP) versus food provisioning (PET), and respective within-group competition for the experimentally provided nut-cracking sites. Previous studies further showed that capuchin monkeys select stones of different sizes and weights to crack different nut species, varying in hardness and size^25,28^.

All animals were tested opportunistically upon group encounter and when they were motivated to participate voluntarily, resulting in a total of 304 trials at SCNP (task 1: N = 117, task 2: N = 122, task 3: N = 65 trials) and 197 trials at PET (task 1: N = 82, task 2: N = 105, task 3: N = 10 trials). At SCNP, 12 individuals (N = 6 males, N = 6 females) out of approximately 40 individuals participated in 2‒51 trials (*M* = 25.33, *SD* = 16.90; SI Table S1). At PET, 15 individuals (N = 7 males, N = 8 females) out of approximately 30 individuals participated in 1‒40 trials (*M* = 11.81, *SD* = 10.87; SI Table S1). All individuals were experienced in nut-cracking using natural stones.

We strictly adhered to the guidelines set out for the ethical treatment of primates established by the American Primatological Society (APS) and the Association for the Study of Animal Behaviour (ASAB). Additionally, the Instituto Brasileiro do Meio Ambiente (IBAMA) and Instituto Chico Mendes de Conservação da Biodiversidade (ICMBio) approved this research (Sisbio authorization 60134).

### 5.2 Experimental set-up

#### Task 1, raw material choice: Do capuchin monkeys discriminate between raw materials?

To investigate stone tool selection according to raw material properties (soft and hard), we provisioned one nut-cracking site with two stones of either raw material and a single nut per trial (Figure 1C, Task 1). This allowed the capuchin monkeys to choose between four stone combinations for nut-cracking (soft anvil & soft hammerstone, soft anvil & hard hammerstone, hard anvil & soft hammerstone, hard anvil & hard hammerstone).

#### Task 2, foraging site selection: Do capuchin monkeys use local enhancement for their selection of a nut-cracking site?

To investigate the role of social cues on the selection of foraging sites, we provisioned two separate nut-cracking sites, each consisting of one stone of either material and a single nut located equidistant between the two sites. At one site, the stones were standardized without percussive damage or evidence of previous use (unmodified stones). At the other site, the stones were artificially modified, and empty nut shells were available (Figure 1D, Task 2). Thus, upon approaching the set-up, capuchin monkeys first needed to select one of the foraging sites and could then choose between the two stone materials. This setup also allowed for the active transportation of stones between each site.

#### Task 3, social cues: Do capuchin monkeys discriminate between stones based on social cues?

To investigate stone tool selection according to asocial and social cues, i.e., raw material properties and signs of previous nut-cracking activities, we modified the set-up of task 1, in that we exchanged one unmodified stone of either material with one modified stone of that material (Figure 1E, Task 3). Thus, capuchin monkeys could select stone tools according to both, the raw materials and the percussive modification.

#### Procedure

Upon group encounter, an area of approximately 16 m² was cleared of all stone or wooden material that could be used for nut-cracking to ensure that only provided stones were used. Additionally, an experimental area of approximately 1 m² was totally cleared of any other material that could distract the monkeys from nut-cracking using the provided stones, leaving an open sandy area. This clearing activity usually attracted individual monkeys to the experimental area so that we could immediately start with testing. To control for potential side or order biases elicited by the experimenter, we set up the stones in a standardized way, moving both hands simultaneously to place two stones at either end of the set up (in a row for Tasks 1 and 3, at individual sides for Task 2). After we started video recording, we placed one nut at an equal distance to all of the four stones to prevent the monkeys to simply select the stones closest to the nut and leave the experimental area and to enable a central approach by the individual most likely to participate next (see also supplementary Video 1).

For all tasks, the stones selected for each trial (presentation of one nut, cashew nut at SCNP, and palm nut at PET) were randomly chosen from the available sample to avoid biases concerning individual stones. The positions of the individual stones (Tasks 1‒3), as well as the foraging sites (Task 2), were randomized between trials to avoid position or side biases. We adjusted the orientation of each trial set-up in the way that the capuchin monkeys would be most likely to approach the set-up centrally to the nut so that all stones were equally likely to be used in terms of their distance and direction to the nut. After each trial, we collected all stones and any nut remains cracked by the subject. For each subsequent trial, we replaced all used stones with new ones to avoid the capuchin monkeys using any potential (odor or visual) cues left behind from the previous use. Before reusing any initially unmodified stones, we removed any traces of nut-cracking activities by abrading their surfaces with sandpaper. If the nut-cracking behavior resulted in heavy or unremovable percussive damage on an unmodified stone, this stone was excluded from all subsequent trials.

Upon group encounter and depending on the stage in testing of the individual approaching the experimental area, we decided which task to set up. For example, if individual A already participated in 10 trials of Task 1, we set up Task 2. When individual A stopped participating and individual B (who has not yet participated in Task 1) approached, we set up Task 1. This flexible set up was possible as all stones were replaced after every single trial and because observers were either subadults (not participating in the experiment) or at further distance (unlikely to view details of the provided stones, thus, reducing a potential observation bias).

To avoid the overrepresentation of specific dominant individuals within one task, additional nut-cracking opportunities were provided away from the experimental task upon need. In doing so, lower-ranking individuals within the groups were provided with the opportunity to participate in the experimental tasks without being chased by more dominant group members.

### 5.3 Data analyses

All experimental trials were videotaped (Sony FDR-AX700) for later behavioral analyses with BORIS (www.boris.unito.it)^74^, starting as soon as the subject picked up the provided nut, or touched the provided stones, and ending when the subject cracked the nut (success) or left the experimental site (failure). For each trial, we recorded the individual IDs of the focal subject (actively exploring the set-up) and of bystanders (observers or scroungers) within a 2 m distance of the experimental set-up. We also noted when the focal was disturbed by any external cues (e.g., group members, other animals), when it used any other objects for nut-cracking than the provided stones, or when it transported the provided stones to other nut-cracking sites to control for potential effects on the nut-cracking performance. These trials (N = 82) were excluded from further statistical analyses. Demographic data was collected in the field for each participating individual, including sex and age class (adult, subadult). Additionally, we recorded each stone ID along with whether it was used as a stone anvil or hammerstone (Table 1bookmark5).

**Table 1.**
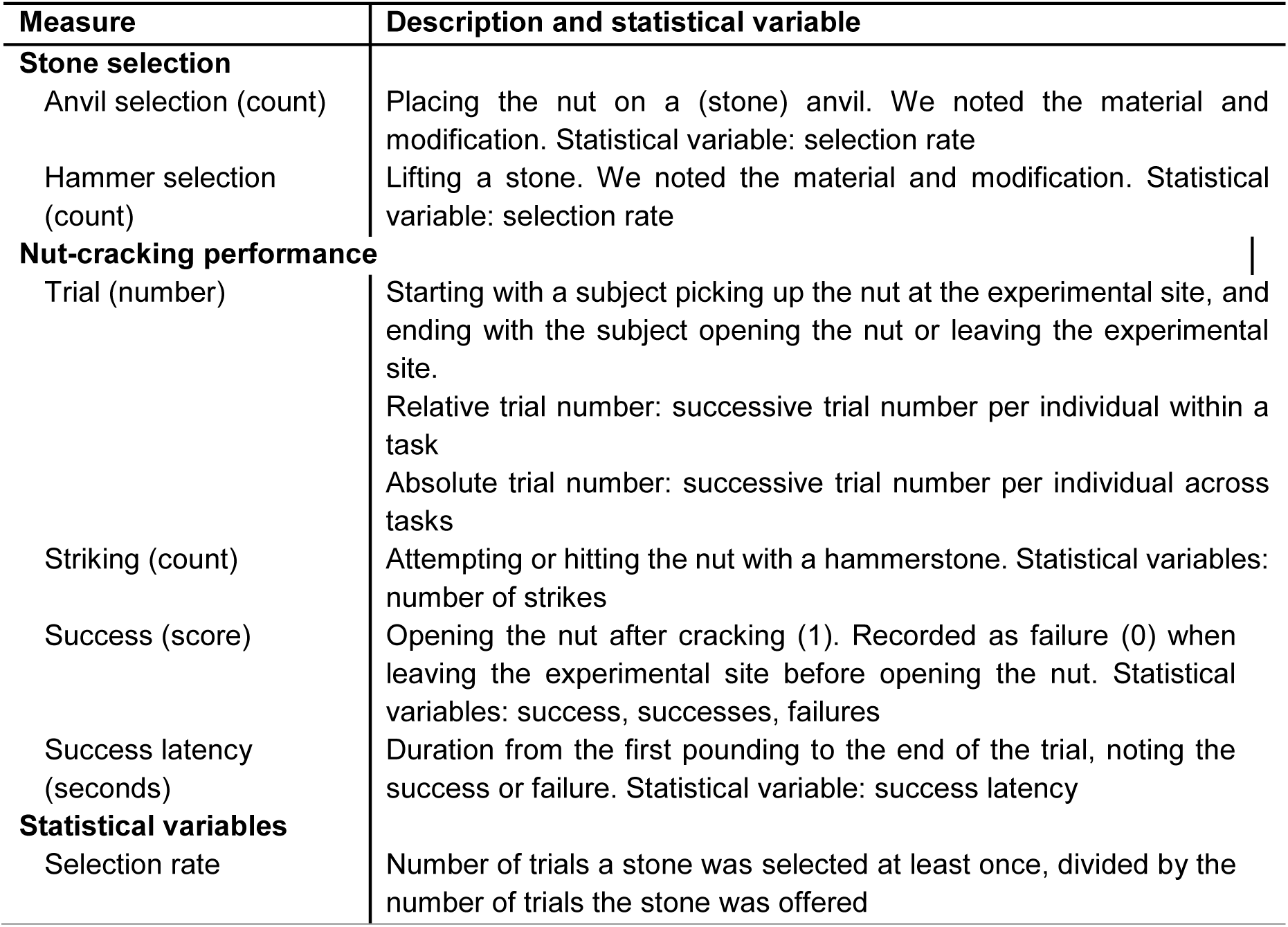
A description of the data associated variables collected from video analyses for each trial.

| Measure | Description and statistical variable |
| --- | --- |
| **Stone selection** |  |
| Anvil selection (count) | Placing the nut on a (stone) anvil. We noted the material and modification. Statistical variable: selection rate |
| Hammer selection (count) | Lifting a stone. We noted the material and modification. Statistical variable: selection rate |
| **Nut-cracking performance** |  |
| Trial (number) | Starting with a subject picking up the nut at the experimental site, and ending with the subject opening the nut or leaving the experimental site.<br>Relative trial number: successive trial number per individual within a task<br>Absolute trial number: successive trial number per individual across tasks |
| Striking (count) | Attempting or hitting the nut with a hammerstone. Statistical variables: number of strikes |
| Success (score) | Opening the nut after cracking (1). Recorded as failure (0) when leaving the experimental site before opening the nut. Statistical variables: success, successes, failures |
| Success latency (seconds) | Duration from the first pounding to the end of the trial, noting the success or failure. Statistical variable: success latency |
| **Statistical variables** |  |
| Selection rate | Number of trials a stone was selected at least once, divided by the number of trials the stone was offered |

### 5.4 Statistical analyses

All statistical analyses were performed using R (version 4.2.3)^75^. We used a series of generalized linear mixed models (GLMM; function “glmer”, package “lme4”)^76^ to examine the effects of our different experimental conditions (see supplementary tables S2-3, S5-10, S12-13 for the individual models). For testing the overall significance of the models, we compared the full models (fixed factors, random slopes, and factors, interactions if applicable) to respective null models, comprising only the intercept and the random effects, using an analysis of variance (ANOVA)^77^. We assessed statistical significance (α-level: 0.5, significance threshold: 0.05) with *Χ²*-tests and calculated the effect sizes (*R²*) of the full models using the function “r.squaredGLMM” (package "MuMIn")^78^ for models with Gaussian error distribution, or calculated Nagelkerke’s *R²* for all other models.

We validated each model and its stability (distribution of random effects, variance of fixed and random effects), and controlled for influential cases by controlling the variance inflation factors (function "vif" applied to standard linear models without any random factors; package "car")^79^, dfbetas (function “dfbeta”), for overdispersion, as well as normality and homogeneity of residuals (qq-plot for the residuals; scatterplot between residuals and fitted values), where applicable. These tests did not reveal any divergence from the model assumptions or influential cases.

#### Task 1, raw material choice: Do capuchin monkeys discriminate between raw materials?

To evaluate stone tool selection according to raw material properties (Figure 1C, Task 1), we investigated the effect of the raw material (limestone/sandstone) on the likelihood that a stone was selected by calculating two separate GLMMs with binomial error structure and a logit-link function (family “binomial”) for either stone anvil or hammerstone selection. We used a binary response of whether a stone was selected (1) or not (0). As fixed effects in the full models, we set the stone material, i.e., either the softer limestone (N = 35) or the harder sandstone (N = 21), the focal’s sex, the study site (PET: N = 12 focals, SCNP: N = 10 focals), the relative trial number (successive trial number per focal within task 1), and as a control the position of the stone. As random factors, we included the stone ID with random slopes for sex, trial, and position, the focal ID with random slopes for material, trial, and position, and the trial ID per focal with a random slope for the material. We excluded correlations between random slopes and intercepts as they seemed unidentifiable (models 1a-b, SI Table S2‒3).

Since the stone anvil and hammerstone selection was dependent on each other, but we investigate their variance in individual models, we ran permutation tests^80,81^ for testing the significance of the full model and the individual predictors. Therefore, we fitted the respective full and null models for N = 999 times with a randomized response variable. To derive the overall significance of the original full model and the individual predictors, we compared the derived test statistics of the randomized data to the test statistics of the original data. Since the stone position encompassed four levels (outer left, centered left, centered right, outer right), we included reduced models lacking the position as a fixed effect in the permutation tests. To estimate the overall significance of the position on the stone selection, we compared the differences in the maximum likelihoods between the reduced models to the respective full models.

#### Task 2, foraging site selection: Do capuchin monkeys use local enhancement for their selection of a nut-cracking site?

To evaluate the role of social cues on the selection of foraging sites (Figure 1D, Task 2), we investigated the effect of the foraging site (unmodified stones versus modified stones and broken nut shells) on the likelihood that a site was selected by calculating a GLMM with binomial error structure and a logit-link function (family “binomial”). We used a binary response of whether a site was selected (1) or not (0). Therefore, we only considered the first site selection within a trial and disregarded when individuals changed the site in the same trial. As fixed effects in the full model, we set the foraging site (unmodified stones/nut-cracking site), the focal’s sex, the study site (PET: N = 13 focals, SCNP: N = 10 focals), the relative trial number (successive trial number per focal within task 2), and as a control the side of the site. As random factors, we included the focal ID with random slopes for trial and position and the trial ID per focal. We excluded correlations between random slopes and intercepts as they seemed unidentifiable (model 2a, SI Table S5).

To evaluate the stone anvil and hammerstone selection between the four different stones (Figure 1D, Task 2), we investigated the effect of the raw material (limestone/sandstone) and signs of previous nut-cracking activities (unmodified stones/modified stones) on the likelihood that a stone was selected. As for task 1, we calculated two separate GLMMs with binomial error structure and a logit-link function (family “binomial”) for either anvil or hammerstone selection. Again, we used a binary response of whether a stone was selected (1) or not (0). As fixed effects in the full models, we set the stone material (limestones: N = 25, sandstones: N = 20), the modification of the stones (unmodified stones: N = 34, modified stones: N = 11), the focal’s sex, the study site (PET: N = 13 focals, SCNP: N = 10 focals), the relative trial number (successive trial number per focal within task 2), and as a control the position of the stone. As random factors, we included the stone ID with random slopes for sex, trial, and position, the focal ID with random slopes for material, stone appearance, trial, and position, and the trial ID per focal with a random slope for material and stone appearance. We excluded correlations between random slopes and intercepts as they seemed unidentifiable (models 2b-c, SI Table S6‒7). Since the stone anvil and hammerstone selection was dependent on each other, but we investigate their variance in individual models, we ran permutation tests similar to task 1 for testing the significance of the full model and the individual predictors. To estimate the overall significance of the position (four levels) on the stone selection, we compared the reduced models lacking the position as a fixed factor to the respective full models.

#### Task 3, social cues: Do capuchin monkeys discriminate between stones based on social cues?

To evaluate stone tool selection according to raw material properties and signs of previous nut-cracking activities (Figure 1E, Task 3), we investigated the effect of the raw material (soft limestone vs. hard sandstone) and signs of previous nut-cracking activities (unmodified stones/modified stones) on the likelihood that a stone was selected. As for tasks 1 and 2, we calculated two separate GLMMs with binomial error structure and a logit-link function (family “binomial”) for either anvil or hammerstone selection. Again, we used a binary response of whether a stone was selected (1) or not (0). As fixed effects in the full models, we set the stone material (limestones: N = 19, sandstones: N = 16), the modification of the stones (unmodified stones: N = 31, modified stones: N = 4), the focal’s sex, the study site (PET: N = 2 focals, SCNP: N = 7 focals), the relative trial number (successive trial number per focal within task 3), and as a control the position of the stone. As random factors, we included the stone ID with random slopes for sex, trial, and position, the focal ID with random slopes for material, stone appearance, trial, and position, and the trial ID per focal with a random slope for material and stone appearance. We excluded correlations between random slopes and intercepts as they seemed unidentifiable (models 3a-b, SI Table S8‒9). Since the anvil and hammerstone selection was dependent on each other, but we investigate their variance in individual models, we ran permutation tests similar to task 1 for testing the significance of the full model and the individual predictors. To estimate the overall significance of the position (four levels) on the stone selection, we compared the reduced models lacking the position as a fixed factor to the respective full models.

#### Nut-cracking performance: Does material choice or individual characteristics influence the nut-cracking performance (over time)?

To evaluate the nut-cracking performance, we first investigated the effect of the last used stone anvil and hammerstone material, the focal’s sex, the study site, and the absolute trial number (successive trial number per focal across tasks) on the success probability. Then, for all successful trials, we evaluated the nut-cracking efficiency, testing the effect of the previous factors on the success latency (nut-cracking duration) and the number of strikes. In all these models, we included random factors: the stone IDs of the stone anvil and hammerstone with random slopes for sex, trial, and the material of the respective stone anvil or hammerstone, the focal ID with random slopes for the stone anvil and hammerstone materials, and trial, and the trial ID per focal, in the case of the success probability and the number of strikes. We excluded correlations between random slopes and intercepts if they seemed unidentifiable.

To evaluate the success probability across trials (tasks 1‒3; PET: N = 189 trials, N = 16 focals; SCNP: N = 304 trials, N = 12 focals), we calculated a GLMM with binomial error structure and a logit-link function (family “binomial”). We used a binary response for the success (1) or failure (0) of nut-cracking per trial (model 4a, SI Table S10).

To evaluate the success latency across successful trials (tasks 1‒3; PET: N = 66 trials, N = 16 focals; SCNP: N = 226 trials, N = 12 focals), we calculated a Gaussian LMM. As the response, we used the log-transformed success latency as the duration a focal needed to crack the nut (model 4b, SI Table S12).

To evaluate the number of strikes across successful trials (tasks 1‒3; PET: N = 66 trials, N = 16 focals; SCNP: N = 226 trials, N = 12 focals), we calculated a Poisson GLMM with log link error function (family = “poisson”). As the response, we used the number of strikes a focal needed to crack the nut (model 4c, SI Table S13).

For all these models, we derived individual test statistics and *p*-values of each predictor by comparing the full model to reduced models, lacking the respective fixed factor, using ANOVA implemented in the function “drop1”.

## Supporting information

Supplementary Model Information

## Acknowledgments

This work was supported by the Max Planck Society and the São Paulo Research Foundation (FAPESP 2018/01292-9; 2019/00716-2). We acknowledge the SCNP and PET management, and the CRAS staff for their permission and support of our data collection, as well as V. H. Tavares de Sousa, A. Macedo, M. Albertin for their assistance in the field. We thank S. Schmid for her assistance in video coding and R. Mundry for statistical advice. The fieldwork was conducted under permit 60134 by Instituto Chico Mendes de Conservação da Biodiversidade (ICMBio) and portaria CNPq N° 846/2022 by the Brazilian Ministry of Science, Technology, and Innovation (MCTI). During writing TP was supported by grant CEECINST/00052/2021 funded by the Portuguese Foundation for Science and Technology (FCT).

## Author Contributions

JHM: conceptualization, study design, data collection, data analysis, manuscript writing, figure production

TF: conceptualization, data collection, manuscript writing

JSR: manuscript writing

HPR: data collection, manuscript writing

TP: study design, data collection, manuscript writing

LVL: project supervision, conceptualization, study design, manuscript writing

