## Supplementary Material for "Social cues on stone tools outweigh raw material properties in wild primates"

*shared first author

Content:

1 Supplementary Results

2 Supplementary Methods

### 1 Supplementary Results

**Table S1** Participation numbers (trials) of 12 wild and 15 semi-wild capuchin monkeys from Serra da Capivara National Park (SCNP) and Park Ecológico do Tietê (PET), respectively. F: female, M: male

| | Task 1 | Task 2 | Task 3 |
|---|---|---|---|
| **SCNP** | 117 | 122 | 65 |
| Assustado (M) | 14 | 20 | 11 |
| Clandestino (M) | 6 | 3 | 4 |
| Encrenquiera (F) | 0 | 7 | 0 |
| Espancado (M) | 8 | 18 | 11 |
| Lica (F) | 15 | 15 | 0 |
| Maca (F) | 15 | 5 | 13 |
| Machucado (M) | 10 | 13 | 10 |
| Maximo (M) | 11 | 19 | 11 |
| Moca (F) | 15 | 13 | 0 |
| Mosca (M) | 1 | 0 | 0 |
| Roma (F) | 3 | 7 | 0 |
| Subadult (F) | 0 | 0 | 5 |
| **PET** | 82 | 105 | 10 |
| Acai (M) | 2 | 3 | 0 |
| Acerola (M) | 5 | 11 | 0 |
| Adult 1 (M) | 1 | 0 | 0 |
| Adult 2 (F) | 14 | 9 | 8 |
| Alex (M) | 1 | 5 | 0 |
| Alice (F) | 1 | 0 | 0 |
| Ana (F) | 0 | 1 | 0 |
| Angelica (F) | 11 | 3 | 1 |
| Caca (F) | 0 | 1 | 0 |
| Caju (M) | 1 | 0 | 0 |
| Capacete (M) | 2 | 1 | 0 |
| Cisca (F) | 1 | 12 | 0 |
| Fisica (F) | 0 | 9 | 0 |
| Frida (F) | 5 | 9 | 0 |
| Frodo (M) | 1 | 4 | 0 |

#### 1.1 Task 1, raw material choice: Do capuchin monkeys discriminate between raw materials?

In addition to the discrimination between stone materials, capuchin monkeys appeared to exhibit a stone anvil selection preference based on stone position (Table S2). Despite the randomized setup, they preferably selected one of the inner stones. Furthermore, individuals at SCNP tended to select fewer stones than individuals at PET, which shows their perseverance for an initially selected stone anvil. Controlling for a potential effect of single individuals driving the model outcome, we compared the full model with a reduced model lacking the random factor “focal ID”, which showed no significance ($X^2(6) = 11.16$, $p = .084$).

**Table S2** Anvil selection: statistical results based on a permutation test (N = 999). Full-null-model comparison: $X^2(4) = 15.03$, $p = .022$; $R^2 = .04$; sd: standard deviation, L: limestone, S: sandstone, F: female, M: male, PET: Park Ecológico do Tietê, SCNP: Serra da Capivara National Park

**model 1a: glmer(selection (yes: 1/no: 0) ~ material + position + sex + park + trial + (1 + position + sex + trial || stone ID) + (1 + material + position + trial || focal ID) + (1 + material || trial ID), family=binomial)**

| **coefficients** | **estimate** | **sd** | ***p*-value** |
|---|---|---|---|
| intercept: soft L, position outer left, sex F, park PET | 0.09 | 0.33 | * |
| material: hard S | –0.64 | 0.24 | 0.002 |
| position | | | 0.001 |
| inner left | 1.15 | 0.32 | * |
| inner right | 0.41 | 0.30 | * |
| outer right | –0.40 | 0.46 | * |
| sex: M | 0.01 | 0.26 | 0.877 |
| park: SCNP | –0.08 | 0.30 | 0.081 |
| trial | –0.09 | 0.11 | 0.247 |

* not shown due to limited interpretation

Similarly, capuchin monkeys appeared to exhibit an additional hammerstone selection preference based on stone position (Table S3), seeming to avoid the outer left stone; and SCNP individuals showed a higher perseverance regarding their hammerstone selection than PET individuals. The permutation test also indicated a significant effect of sex, with males showing a lower perseverance than females, and of trial, indicating a decreasing perseverance throughout experimental participation. While the effect of sex may arise due to the unequally distributed participation rates between females and males, the effect of trial may hint towards more complex influences based on motivation and experience. For instance, subjects learned over the course of the trials that they were safe to participate at the experiment, having the time to switch between stones to find a more efficient tool. Controlling for a potential effect of single individuals driving the model outcome, we compared the full model with a reduced model lacking the random factor “focal ID”, which showed no significance ($X^2(6) = 7.19$, $p = .304$).

**Table S3** Hammer selection: statistical results based on a permutation test (N = 999). Full-null-model comparison: $X^2(4) = 1.56$, $p = .004$; $R^2 = .02$; sd: standard deviation, L: limestone, S: sandstone, F: female, M: male, PET: Park Ecológico do Tietê, SCNP: Serra da Capivara National Park

**model 1b: glmer(selection (yes: 1/no: 0) ~ material + position + sex + park + trial + (1 + position + sex + trial || stone ID) + (1 + material + position + trial || focal ID) + (1 + material || trial ID), family=binomial)**

| **coefficients** | **estimate** | **sd** | ***p*-value** |
|---|---|---|---|
| intercept: soft L, position outer left, sex F, park PET | –1.22 | 0.37 | * |
| material: hard S | 0.82 | 0.32 | 0.001 |
| position | | | 0.018 |
| inner left | –0.32 | 0.44 | * |
| inner right | 0.14 | 0.33 | * |

| | | | |
|---|---|---|---|
| outer right | –1.28 | 0.41 | * |
| sex: M | 0.18 | 0.25 | 0.001 |
| park: SCNP | –0.28 | 0.30 | 0.014 |
| trial | 0.05 | 0.12 | 0.004 |

* not shown due to limited interpretation

#### 1.2 Tool material choice

**Table S4** Number of times each stone material was chosen as anvil or hammer across the three tasks. Since individuals changed anvils or hammers within trials, the numbers do not necessarily sum up to the total trial number per task.

| | Soft anvil | Hard anvil | Soft hammer | Hard hammer |
|---|---|---|---|---|
| **Task 1 (152 trials)** | 130 | 98 | 67 | 103 |
| **Task 2 (189 trials)** | 179 | 120 | 93 | 127 |
| **Task 3 (73 trials)** | 69 | 54 | 43 | 52 |

#### 1.3 Task 2, foraging site selection: Do capuchin monkeys use local enhancement for their selection of a nut-cracking site?

Controlling for a potential effect of single individuals driving the model outcome, we compared the full model with a reduced model lacking the random factor “focal ID”, which was significant ($X^2(4) = 77.14$, $p < .001$), but the visual inspection shows that most of the individuals chose the nut-cracking site (modified stones) more frequently than the foraging site with unmodified stones (not shown), confirming the results of model 2a (Table S5).

**Table S5** Foraging-site selection: statistical results based on full-reduced-model comparisons (drop1). Full-null-model comparison: $X^2(4) = 14.19$, $p = .007$; $R^2 = .05$; sd: standard deviation, F: female, M: male, PET: Park Ecológico do Tietê, SCNP: Serra da Capivara National Park

| **model 2a: glmer(selection (yes: 1/no: 0) ~ foraging site + side + sex + park + trial + (1 + site + side + trial \|\| focal ID) + (1 \| trial ID), family=binomial)** | | | |
|---|---|---|---|
| **coefficients** | **estimate** | **sd** | ***p*-value** |
| intercept: modified stones, side left, sex F, park PET | 1.02 | 0.44 | * |
| foraging site: unmodified stones | –1.86 | 0.40 | <0.001 |
| side: right | –0.17 | 0.64 | 0.790 |
| sex: M | 0.00 | 0.24 | 1.000 |
| park: SCNP | 0.00 | 0.24 | 1.000 |
| trial | 0.00 | 0.03 | 1.000 |

* not shown due to limited interpretation

In addition to the significant effects of stone material and modification on stone anvil selection, SCNP individuals showed a higher perseverance regarding their anvil selection than PET individuals, while males showed a lower perseverance than females (Table S5). Further, the perseverance for a specific stone selection tended to increase throughout experimental participation. Controlling for a potential effect of single individuals driving the model outcome, we compared the full model with a reduced model lacking the random factor “focal ID”, which was significant ($X^2(7) = 32.75$, $p < .001$), but visual inspections show that most of the individuals chose the limestones over sandstones (not shown) and modified over unmodified stones (not shown), confirming the results of model 2b (Table S6).

**Table S6** Anvil selection: statistical results based on a permutation test (N = 999). Full-null-model comparison: $X^2(5) = 23.87$, $p = .001$; $R^2 = .08$; sd: standard deviation, L: limestone, S: sandstone, F: female, M: male, PET: Park Ecológico do Tietê, SCNP: Serra da Capivara National Park

**model 2b: glmer(selection (yes: 1/no: 0) ~ material + modification + position + sex + park + trial + (1 + position + sex + trial || stone ID) + (1 + material + modification + position + trial||ID) + (1 + material + modification || trial ID), family=binomial)**

| **coefficients** | **estimate** | **sd** | ***p*-value** |
|---|---|---|---|
| intercept: soft L, unmodified, position outer left, sex F, park PET | –0.20 | 0.30 | * |
| material: hard S | –0.91 | 0.20 | 0.001 |
| modification: modified | 1.21 | 0.31 | 0.001 |
| position | | | 0.376 |
| inner left | 0.18 | 0.25 | * |
| inner right | 0.51 | 0.39 | * |
| outer right | 0.02 | 0.37 | * |
| sex: M | 0.12 | 0.22 | 0.018 |
| park: SCNP | –1.19 | 0.21 | 0.001 |
| trial | –0.08 | 0.09 | 0.090 |

* not shown due to limited interpretation

Likewise, SCNP individuals showed a higher perseverance than PET individuals regarding their hammerstone selection in Task 2 (Table S6). Controlling for a potential effect of single individuals driving the model outcome, we compared the full model with a reduced model lacking the random factor “focal ID”, which was significant ($X^2(7) = 50.25$, $p < .001$), but visual inspections show that most of the individuals chose the sandstones over limestones (not shown) and modified over unmodified stones (not shown), confirming the results of model 2c (Table S7).

**Table S7** Hammer selection: statistical results based on a permutation test (N = 999). Full-null-model comparison: $X^2(5) = 6.81$, $p = .016$; $R^2 = .03$; sd: standard deviation, L: limestone, S: sandstone, F: female, M: male, PET: Park Ecológico do Tietê, SCNP: Serra da Capivara National Park

**model 2c: glmer(selection (yes: 1/no: 0) ~ material + modification + position + sex + park + trial + (1 + position + sex + trial || stone ID) + (1 + material + modification + position + trial||ID) + (1 + material + modification || trial ID), family=binomial)**

| **coefficients** | **estimate** | **sd** | ***p*-value** |
|---|---|---|---|
| intercept: soft L, unmodified, position outer left, sex F, park PET | –2.02 | 0.48 | * |
| material: hard S | 0.68 | 0.34 | 0.022 |
| modification: modified | 1.27 | 0.42 | 0.002 |
| position | | | 0.376 |
| inner left | –0.39 | 0.50 | * |

| | | | |
|---|---|---|---|
| inner right | –0.65 | 0.49 | * |
| outer right | 0.42 | 0.54 | * |
| sex: M | 0.05 | 0.25 | 0.763 |
| park: SCNP | –0.52 | 0.28 | 0.017 |
| trial | –0.01 | 0.12 | 0.450 |

* not shown due to limited interpretation

#### 1.4 Task 3, social cues: Do capuchin monkeys discriminate between stones based on social cues?

Controlling for a potential effect of single individuals driving the model outcome, we compared the full model with a reduced model lacking the random factor “focal ID”, which was significant ($X^2(7) = 20.49$, $p = .005$), and plotted the stone selection of each individuals, validating that none of the individuals was driving the model output but most of the individuals chose modified over unmodified stones (not shown), confirming the results of model 3a (Table S8).

**Table S8** Anvil selection: statistical results based on a permutation test (N = 999). Full-null-model comparison: $X^2(5) = 5.00$, $p = .004$; $R^2 = .07$; sd: standard deviation, L: limestone, S: sandstone, F: female, M: male, PET: Park Ecológico do Tietê, SCNP: Serra da Capivara National Park

**model 3a: glmer(selection (yes: 1/no: 0) ~ material + modification + position + sex + park + trial + (1 + position + sex + trial || stone ID) + (1 + material + modification + position + trial || focal ID) + (1 + material + modification || trial ID), family=binomial)**

| **coefficients** | **estimate** | **sd** | ***p*-value** |
|---|---|---|---|
| intercept: soft L, unmodified, position outer left, sex F, park PET | 0.50 | 0.66 | * |
| material: hard S | –0.37 | 0.35 | 0.255 |
| modification: modified | 1.08 | 0.30 | 0.003 |
| position | | | 0.230 |
| inner left | –0.50 | 0.77 | * |
| inner right | –0.86 | 0.69 | * |
| outer right | –1.12 | 0.52 | * |
| sex: M | 0.38 | 0.42 | 0.257 |
| park: SCNP | –1.01 | 0.63 | 0.106 |
| trial | 0.12 | 0.14 | 0.466 |

* not shown due to limited interpretation

Controlling for a potential effect of single individuals driving the model outcome, we compared the full model with a reduced model lacking the random factor “focal ID”, which showed no significance ($X^2(7) = 13.28$, $p = .066$).

**Table S9** Hammer selection: statistical results based on a permutation test (N = 999). Full-null-model comparison: $X^2(5) = 2.63$, $p = .107$; $R^2 = .02$; sd: standard deviation, L: limestone, S: sandstone, F: female, M: male, PET: Park Ecológico do Tietê, SCNP: Serra da Capivara National Park

**model 3b: glmer(selection (yes: 1/no: 0) ~ material + modification + position + sex + park + trial + (1 + position + sex + trial || stone ID) + (1 + material + modification + position + trial || focal ID) + (1 + material + modification || trial ID), family=binomial)**

| coefficients | estimate | sd | *p*-value |
|---|---|---|---|
| intercept: soft L, unmodified, position outer left, sex F, park PET | –2.09 | 0.84 | * |
| material: hard S | 0.50 | 0.37 | 0.247 |
| modification: modified | –0.65 | 0.39 | 0.112 |
| position | | | 0.213 |
| inner left | 1.60 | 0.80 | * |
| inner right | 0.96 | 0.74 | * |
| outer right | 0.36 | 0.79 | * |
| sex: M | 0.17 | 0.37 | 0.669 |
| park: SCNP | 0.22 | 0.66 | 0.057 |
| trial | –0.03 | 0.16 | 0.346 |

* not shown due to limited interpretation

#### 1.5 Nut-cracking performance and efficiency

Controlling for a potential effect of single individuals driving the model outcome for the probability to successfully crack the provided nuts, we compared the full model with a reduced model lacking the random factor “focal ID”, which showed no significance ($X^2(4) = 3.44$, $p = .487$), suggesting equal nut-cracking proficiency across individuals.

Table S10 Nut-cracking success: statistical results based on full-reduced-model comparisons (drop1). Full-null-model comparison: $X^2(5) = 22.28$, $p < .001$; $R^2 = .08$; sd: standard deviation, L: limestone, S: sandstone, F: female, M: male, PET: Park Ecológico do Tietê, SCNP: Serra da Capivara National Park

**model 4a: glmer(success (yes: 1/no: 0) ~ anvil material + hammer material + trial + sex + park + (1 + hammer material + trial + sex | stone anvil ID) + (1 + anvil material + trial + sex | hammerstone ID) + (1 + hammer material + anvil material + trial | focal ID) + (1 | trial ID), family=binomial, control= glmerControl(optimizer="bobyqa", optCtrl=list(maxfun=100000))))**

| coefficients | estimate | sd | *p*-value |
|---|---|---|---|
| intercept: anvil: soft L, hammer: soft L, sex F, park PET | 0.09 | 0.47 | * |
| anvil material: hard S | 0.04 | 0.40 | 0.920 |
| hammer material: hard S | 0.16 | 0.34 | 0.632 |
| sex: M | –0.25 | 0.40 | 0.521 |
| park: SCNP | 1.54 | 0.43 | <0.001 |
| trial | 0.51 | 0.17 | 0.005 |

* not shown due to limited interpretation

**Table S11** Failures to crack the provided nut using the provided stones at Park Ecológico do Tietê (PET) and Serra da Capivara National Park (SCNP)

| reason of failure | task | PET | SCNP |
|---|---|---|---|
| losing nut | 1 | 21 | 4 |
| | 2 | 14 | 10 |

| | | | |
|---|---|---|---|
| | 3 | 0 | 0 |
| giving-up | 1 | 8 | 0 |
| | 2 | 4 | 5 |
| | 3 | 0 | 3 |
| leaving with nut | 1 | 0 | 11 |
| | 2 | 1 | 4 |
| | 3 | 0 | 0 |
| discarded nut | 1 | 2 | 2 |
| | 2 | 4 | 0 |
| | 3 | 0 | 0 |
| being chased | 1 | 1 | 1 |
| | 2 | 4 | 5 |
| | 3 | 1 | 2 |

Controlling for a potential effect of single individuals driving the model outcome for the success latency, we compared the full model with a reduced model lacking the random factor "focal ID", which was significant ($X^2(10) = 37.61$, $p < .001$), showing individual variation in nut-cracking speed.

Table S12 Nut-cracking efficiency, success latency: statistical results based on full-reduced-model comparisons (drop1). Full-null-model comparison: $X^2(5) = 2.16$, $p = .826$; $R^2 = .34$; sd: standard deviation, L: limestone, S: sandstone, F: female, M: male, PET: Park Ecológico do Tietê, SCNP: Serra da Capivara National Park

| **model 4b: lmer(log-transformed success latency ~ anvil material + hammer material + trial + sex + park + (1 + hammer material + trial + sex \|\| stone anvil ID) + (1 + anvil material + trial + sex \|\| hammerstone ID) + (1 + hammer material + anvil material + trial \|\| focal ID), control= lmerControl(optimizer="bobyqa", optCtrl=list(maxfun=100000)))** | | | |
|---|---|---|---|
| **coefficients** | **estimate** | **sd** | ***p*-value** |
| intercept: anvil: soft L, hammer: soft L, sex F, park PET | 2.61 | 0.26 | * |
| anvil material: hard S | 0.21 | 0.15 | 0.189 |
| hammer material: hard S | 0.01 | 0.12 | 0.965 |
| sex: M | –0.01 | 0.24 | 0.980 |
| park: SCNP | –0.10 | 0.25 | 0.749 |
| trial | –0.03 | 0.08 | 0.659 |

* not shown due to limited interpretation

Controlling for a potential effect of single individuals driving the model outcome for the nut-cracking efficiency in terms of the number of strikes until nut-cracking, we compared the full model with a reduced model lacking the random factor "focal ID", which was significant ($X^2(10) = 41.61$, $p < .001$), showing individual variation in nut-cracking efficiency.

**Table S13** Nut-cracking efficiency, number of strikes: statistical results based on full-reduced-model comparisons (drop1). Full-null-model comparison: $X^2(5) = 4.18$, $p = .525$; $R^2 = .01$; sd: standard deviation, L: limestone, S: sandstone, F: female, M: male, PET: Park Ecológico do Tietê, SCNP: Serra da Capivara National Park

**model 4c: glmer(number of strikes ~ anvil material + hammer material + trial + sex + park + (1 + hammer material + trial + sex || stone anvil ID) + (1 + anvil material + trial + sex || hammerstone ID) + (1 + hammer material + anvil material + trial || focal ID) + (1 | trial ID), family=poisson, control=lmerControl(optimizer="bobyqa", optCtrl=list(maxfun=100000)))**

| **coefficients** | **estimate** | **sd** | ***p*-value** |
|---|---|---|---|
| intercept: anvil: soft L, hammer: soft L, sex F, park PET | 1.25 | 0.20 | * |
| anvil material: hard S | 0.08 | 0.12 | 0.518 |
| hammer material: hard S | –0.01 | 0.12 | 0.965 |
| sex: M | 0.08 | 0.18 | 0.682 |
| park: SCNP | 0.31 | 0.20 | 0.125 |
| trial | 0.03 | 0.06 | 0.625 |

* not shown due to limited interpretation

### 2 Supplementary Methods

#### 2.1 Determining natural raw material properties of capuchin nut cracking tools and preparation of experimental hammerstones and anvils

To assess the relative hardness of capuchin anvils and hammerstones in a wild setting, we sampled natural or habitually used nut-cracking sites at both SCNP and PET, respectively. We measured the elastic rebound hardness of a representative sample of the same raw material types used as both hammerstones and anvils using a Schmidt hammer. Schmidt hammer values are recorded on a relative scale between 0 to 100 with increasing hardness, i.e., decreasing elasticity. Since Schmidt hammer values vary with stone size and movement[57], all measured samples were similar-sized, embedded stones. For each sample, ten rebound hardness values were taken which were averaged to provide a mean hardness value for each raw material.

We found a significant difference in the hardness of stone anvils and hammerstones used in the wild at SCNP, with hammerstones being considerably harder compared to anvils (t-test: $t(51) = 22.63$, $p < .001$). At SCNP, hammerstones are usually hard quartzite cobbles, with a mean rebound hardness value of 69.79 ± 6.96 (N = 35), and anvils are usually softer sandstone blocks, with a mean rebound hardness of 29.35 ± 6.30 (N = 23). Conversely at PET, there is no significant difference in the hardness of stones used as hammers and anvils (t-test: $t(6) = 2.01$, $p = .090$). At PET, hammerstones are mostly granite blocks, with a mean rebound hardness of 32.16 ± 6.56 (N = 7), and anvils are mostly concrete pavements, with a mean rebound hardness of 37.67 ± 0.47 (N = 3).

Based on the hardness of natural stone tools of wild capuchin monkeys (SCNP), we selected one softer (fine-grained limestone; rebound hardness mean: 27.05 ± 11.28; N = 69) and one harder stone material (metamorphosed sandstone; rebound hardness mean: 57.09 ± 8.39; $N = 37$), significantly differing in their relative hardness (t-test: $t(29.8) = -15.36$, $p < .001$; **Fig. S1**). The rebound hardness of these materials was measured on a small sample of the materials to not inflict any damage by the Schmidt hammer on the actual experimental stones (Table S13).

The dimensions of all experimental stones were standardized to equally sized cuboids (6 cm x 6 cm x 4 cm). Both raw material types possessed a comparable. Due to the varying densities of the two materials, the harder sandstones were slightly heavier (mean = 360.30 ± 6.35 g, $N = 25$) than the softer limestones (mean = 322.49 ± 34.34 g, $N = 51$).

For the modified stones, we artificially produced signs of previous nut-cracking activities (social cues) on a subset of these standardized stones by inflicting typical percussive damage as well as adding visible nut-shell residue. To mimic natural hammerstone damage, pieces of the harder sandstone were percussed superficially along their edges without severe pits or detachments. To mimic natural stone anvil damage, pieces of the softer limestone were percussed both along their edges as well as on a single active surface, resulting in a combination of percussive damage along the edges and a notable central pitted region. In addition to this physical damage, nut residue was adhered to the entire surface of the presumed sandstone hammers and predominantly to the upper active surface of the presumed limestone anvils.

#### 2.2 Sampling of natural and experimental stones

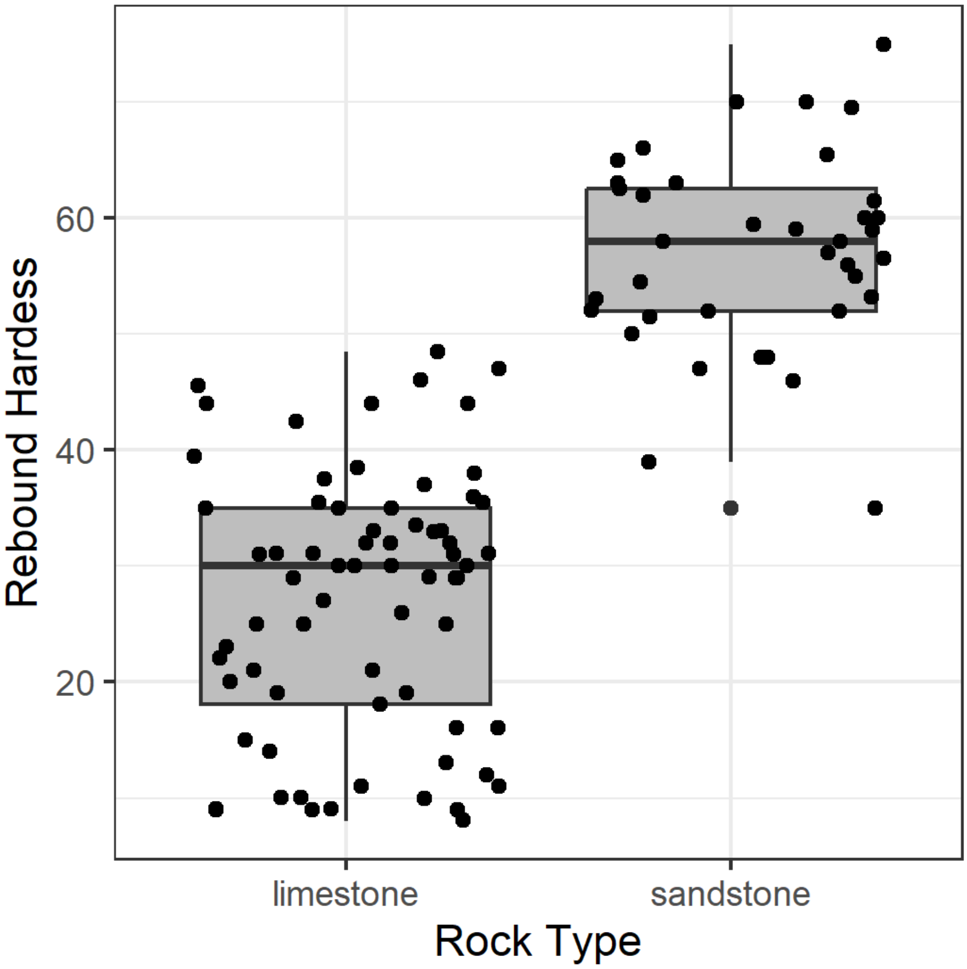


**Fig. S1** Rebound hardness measures for N = 69 experimental limestones and N = 37 experimental sandstones (standardized dimensions: 6 x 6 x 4 cm). Limestones (mean = 27.1 ± 11.9) were significantly softer than sandstones (mean = 57.1 ± 8.4; $t(29.8) = -15.36$, $p < 0.001$)

**Table S14** Rebound hardness measures (averages over 10 measures) for the sampled natural and experimental stones. Shown are individual measures, as well as mean and standard deviation (sd) over environments and materials. SCNP: Serra da Capivara National Park, PET: Park Ecológico do Tietê

| **sample ID** | **environment** | **raw material** | size | mass | **rebound hardness** | **N** |
|---|---|---|---|---|---|---|
| 53 | SCNP | concrete | | | 33.0 | **1** |
| | **SCNP** | **quartzite** | | | **mean = 69.8±7.0** | **35** |
| 10 | SCNP | quartzite | | | 80.0 | |
| 21 | SCNP | quartzite | | | 72.0 | |
| 22 | SCNP | quartzite | | | 56.5 | |
| 23 | SCNP | quartzite | | | 76.0 | |
| 24 | SCNP | quartzite | | | 63.5 | |
| 25 | SCNP | quartzite | | | 79.0 | |
| 26 | SCNP | quartzite | | | 75.5 | |
| 27 | SCNP | quartzite | | | 75.0 | |
| 28 | SCNP | quartzite | | | 67.0 | |
| 29 | SCNP | quartzite | | | 71.0 | |
| 30 | SCNP | quartzite | | | 68.0 | |

| | | | | |
|---|---|---|---|---|
| 31 | SCNP | quartzite | 74.0 | |
| 32 | SCNP | quartzite | 76.5 | |
| 33 | SCNP | quartzite | 76.5 | |
| 34 | SCNP | quartzite | 73.5 | |
| 35 | SCNP | quartzite | 77.0 | |
| 36 | SCNP | quartzite | 67.5 | |
| 37 | SCNP | quartzite | 66.5 | |
| 38 | SCNP | quartzite | 66.0 | |
| 39 | SCNP | quartzite | 78.0 | |
| 40 | SCNP | quartzite | 76.0 | |
| 48 | SCNP | quartzite | 65.0 | |
| 49 | SCNP | quartzite | 59.0 | |
| 51 | SCNP | quartzite | 75.5 | |
| 52 | SCNP | quartzite | 53.0 | |
| 54 | SCNP | quartzite | 54.0 | |
| 57 | SCNP | quartzite | 72.5 | |
| 58 | SCNP | quartzite | 64.5 | |
| 59 | SCNP | quartzite | 69.0 | |
| 60 | SCNP | quartzite | 71.0 | |
| 62 | SCNP | quartzite | 70.5 | |
| 63 | SCNP | quartzite | 77.0 | |
| 64 | SCNP | quartzite | 64.5 | |
| 65 | SCNP | quartzite | 68.5 | |
| 66 | SCNP | quartzite | 63.5 | |
| | **SCNP** | **sandstone** | **mean = 29.4±6.3** | **23** |
| 7 | SCNP | sandstone | 26.0 | |
| 8 | SCNP | sandstone | 31.5 | |
| 9 | SCNP | sandstone | 27.0 | |
| 11 | SCNP | sandstone | 28.5 | |
| 12 | SCNP | sandstone | 29.5 | |
| 13 | SCNP | sandstone | 35.0 | |
| 14 | SCNP | sandstone | 32.5 | |
| 15 | SCNP | sandstone | 25.5 | |
| 16 | SCNP | sandstone | 21.0 | |
| 17 | SCNP | sandstone | 29.5 | |
| 18 | SCNP | sandstone | 28.0 | |
| 19 | SCNP | sandstone | 20.5 | |
| 20 | SCNP | sandstone | 29.0 | |
| 41 | SCNP | sandstone | 36.0 | |
| 42 | SCNP | sandstone | 33.0 | |
| 43 | SCNP | sandstone | 41.5 | |
| 44 | SCNP | sandstone | 38.0 | |
| 45 | SCNP | sandstone | 28.5 | |
| 46 | SCNP | sandstone | 23.5 | |
| 47 | SCNP | sandstone | 16.5 | |
| 55 | SCNP | sandstone | 25.0 | |
| 56 | SCNP | sandstone | 32.0 | |
| 61 | SCNP | sandstone | 41.5 | |
| | **SCNP** | **wood** | **mean = 25.3±11.2** | **6** |
| 1 | SCNP | wood | 41.0 | |
| 2 | SCNP | wood | 38.0 | |
| 3 | SCNP | wood | 19.0 | |
| 4 | SCNP | wood | 26.0 | |

| | | | | | | |
|---|---|---|---|---|---|---|
| 5 | SCNP | wood | | | 19.0 | |
| 6 | SCNP | wood | | | 9.0 | |
| | **PET** | **concrete** | | | **mean = 37.7±0.5** | **3** |
| 69 | PET | concrete | | | 38.0 | |
| 71 | PET | concrete | | | 38.0 | |
| 76 | PET | concrete | | | 37.0 | |
| | **PET** | **granite** | | | **mean = 32.2±6.6** | **7** |
| 67 | PET | granite | | | 40.0 | |
| 68 | PET | granite | | | 42.5 | |
| 70 | PET | granite | | | 26.5 | |
| 72 | PET | granite | | | 26.5 | |
| 73 | PET | granite | | | 34.5 | |
| 74 | PET | granite | | | 25.0 | |
| 77 | PET | granite | | | 31.5 | |
| | **PET** | **wood** | | | **mean = 35.3±4.2** | **2** |
| 75 | PET | wood | | | 31.5 | |
| 78 | PET | wood | | | 39.5 | |
| | **experiment** | **limestone** | **6x6x4 cm** | **mean = 322.5±34.3 (N=51)** | **mean = 27.1±11.9 (N=69)** | **71** |
| 117 | experiment | limestone | | | 29.0 | |
| 118 | experiment | limestone | | | 21.0 | |
| 119 | experiment | limestone | | | 14.0 | |
| 120 | experiment | limestone | | | 26.0 | |
| 121 | experiment | limestone | | | 20.5 | |
| 122 | experiment | limestone | | | 10.5 | |
| 123 | experiment | limestone | | | 11.5 | |
| 124 | experiment | limestone | | | 19.5 | |
| 125 | experiment | limestone | | | 16.5 | |
| 126 | experiment | limestone | | | 13.0 | |
| 127 | experiment | limestone | | | 18.5 | |
| 128 | experiment | limestone | | | 10.5 | |
| 129 | experiment | limestone | | | 9.0 | |
| 130 | experiment | limestone | | | 9.0 | |
| 131 | experiment | limestone | | | 9.0 | |
| 132 | experiment | limestone | | | 10.0 | |
| 133 | experiment | limestone | | | 11.0 | |
| 134 | experiment | limestone | | | 8.5 | |
| 135 | experiment | limestone | | | 9.5 | |
| 136 | experiment | limestone | | | 9.5 | |
| L2 | **experiment** | **limestone** | | **320.4** | **29.5** | |
| L3 | **experiment** | **limestone** | | **334.2** | **31.0** | |
| L6 | **experiment** | **limestone** | | **332.9** | **32.0** | |
| L14 | **experiment** | **limestone** | | **518.3** | **23.5** | |
| L17 | **experiment** | **limestone** | | **415.2** | **19.5** | |
| L114 | **experiment** | **limestone** | | **313.5** | **38.5** | |
| L115 | **experiment** | **limestone** | | **327.0** | **39.5** | |
| L118 | **experiment** | **limestone** | | **310.0** | **33.0** | |
| L119 | **experiment** | **limestone** | | **280.3** | **44.0** | |
| L120 | **experiment** | **limestone** | | **326.6** | **31.5** | |
| L123 | **experiment** | **limestone** | | **291.4** | **22.5** | |
| L124 | **experiment** | **limestone** | | **308.2** | **12.0** | |
| L125 | **experiment** | **limestone** | | **326.3** | **44.0** | |

| | | | | | | |
|---|---|---|---|---|---|---|
| L126 | **experiment** | **limestone** | | **305.2** | **16.5** | |
| L180 | **experiment** | **limestone** | | **324.0** | **33.5** | |
| L132 | **experiment** | **limestone** | | **316.5** | **31.5** | |
| L134 | **experiment** | **limestone** | | **273.9** | **27.0** | |
| L135 | **experiment** | **limestone** | | **311.0** | **37.0** | |
| L136 | **experiment** | **limestone** | | **317.4** | **33.0** | |
| L137 | **experiment** | **limestone** | | **282.3** | **46.0** | |
| L140 | **experiment** | **limestone** | | **328.1** | **30.5** | |
| L143 | **experiment** | **limestone** | | **296.2** | **30.5** | |
| L146 | **experiment** | **limestone** | | **320.7** | **35.0** | |
| L149 | **experiment** | **limestone** | | **292.8** | **29.0** | |
| L151 | **experiment** | **limestone** | | **315.8** | **35.5** | |
| L152 | **experiment** | **limestone** | | **325.1** | **35.5** | |
| L153 | **experiment** | **limestone** | | **312.9** | **37.5** | |
| L154 | **experiment** | **limestone** | | **320.2** | **31.5** | |
| L156 | **experiment** | **limestone** | | **335.8** | **44.0** | |
| L158 | **experiment** | **limestone** | | **318.5** | **35.0** | |
| L159 | **experiment** | **limestone** | | **335.1** | **21.0** | |
| L160 | **experiment** | **limestone** | | **320.9** | **36.0** | |
| L161 | **experiment** | **limestone** | | **316.0** | **15.0** | |
| L181 | **experiment** | **limestone** | | **309.0** | **30.5** | |
| L162 | **experiment** | **limestone** | | **321.2** | **38.0** | |
| L163 | **experiment** | **limestone** | | **323.7** | **31.5** | |
| L164 | **experiment** | **limestone** | | **290.7** | **30.0** | |
| L165 | **experiment** | **limestone** | | **318.0** | **25.0** | |
| L166 | **experiment** | **limestone** | | **338.7** | **48.5** | |
| L167 | **experiment** | **limestone** | | **327.2** | **47.0** | |
| L168 | **experiment** | **limestone** | | **327.5** | **33.0** | |
| L169 | **experiment** | **limestone** | | **310.3** | **29.0** | |
| L171 | **experiment** | **limestone** | | **334.9** | **42.5** | |
| L172 | **experiment** | **limestone** | | **330.5** | **45.5** | |
| L173 | **experiment** | **limestone** | | **301.0** | **25.0** | |
| L174 | **experiment** | **limestone** | | **302.7** | **25.0** | |
| L175 | **experiment** | **limestone** | | **318.0** | **35.0** | |
| L177 | **experiment** | **limestone** | | **328.4** | **32.0** | |
| L180 | **experiment** | **limestone** | | **324.0** | **32.0** | |
| L90 | **experiment** | **limestone** | | **337.0** | **NA** | |
| L92 | **experiment** | **limestone** | | **331.6** | **NA** | |
| | **experiment** | **sandstone** | **6x6x4 cm** | **mean = 360.3±6.4 (N=25)** | **mean = 57.1±8.4** | **37** |
| 107 | experiment | sandstone | | | 55.0 | |
| 108 | experiment | sandstone | | | 65.5 | |
| 109 | experiment | sandstone | | | 61.5 | |
| 110 | experiment | sandstone | | | 59.0 | |
| 111 | experiment | sandstone | | | 62.0 | |
| 112 | experiment | sandstone | | | 66.0 | |
| 113 | experiment | sandstone | | | 59.5 | |
| 114 | experiment | sandstone | | | 62.5 | |
| 115 | experiment | sandstone | | | 60.0 | |
| 116 | experiment | sandstone | | | 63.0 | |

| | | | | |
|---|---|---|---|---|
| 137 | experiment | sandstone | | 56.0 |
| 138 | experiment | sandstone | | 53.2 |
| S44 | **experiment** | **sandstone** | **357.7** | **52.0** |
| S45 | **experiment** | **sandstone** | **355.2** | **65.0** |
| S46 | **experiment** | **sandstone** | **370.0** | **46.0** |
| S47 | **experiment** | **sandstone** | **357.7** | **54.5** |
| S48 | **experiment** | **sandstone** | **358.6** | **51.5** |
| S49 | **experiment** | **sandstone** | **358.5** | **58.0** |
| S50 | **experiment** | **sandstone** | **360.0** | **59.0** |
| S51 | **experiment** | **sandstone** | **357.9** | **70.0** |
| S91 | **experiment** | **sandstone** | **371.4** | **39.0** |
| S52 | **experiment** | **sandstone** | **362.6** | **52.0** |
| S53 | **experiment** | **sandstone** | **359.1** | **52.0** |
| S54 | **experiment** | **sandstone** | **351.8** | **47.0** |
| S68 | **experiment** | **sandstone** | **359.0** | **35.0** |
| S75 | **experiment** | **sandstone** | **372.2** | **75.0** |
| S76 | **experiment** | **sandstone** | **354.0** | **56.5** |
| S77 | **experiment** | **sandstone** | **362.3** | **48.0** |
| S78 | **experiment** | **sandstone** | **358.2** | **48.0** |
| S79 | **experiment** | **sandstone** | **346.2** | **57.0** |
| S80 | **experiment** | **sandstone** | **362.8** | **70.0** |
| S81 | **experiment** | **sandstone** | **358.4** | **53.0** |
| S82 | **experiment** | **sandstone** | **358.1** | **60.0** |
| S83 | **experiment** | **sandstone** | **365.1** | **58.0** |
| S84 | **experiment** | **sandstone** | **368.6** | **50.0** |
| S85 | **experiment** | **sandstone** | **352.1** | **63.0** |
| S86 | **experiment** | **sandstone** | **370.1** | **69.5** |

### 3 Supplementary Figures

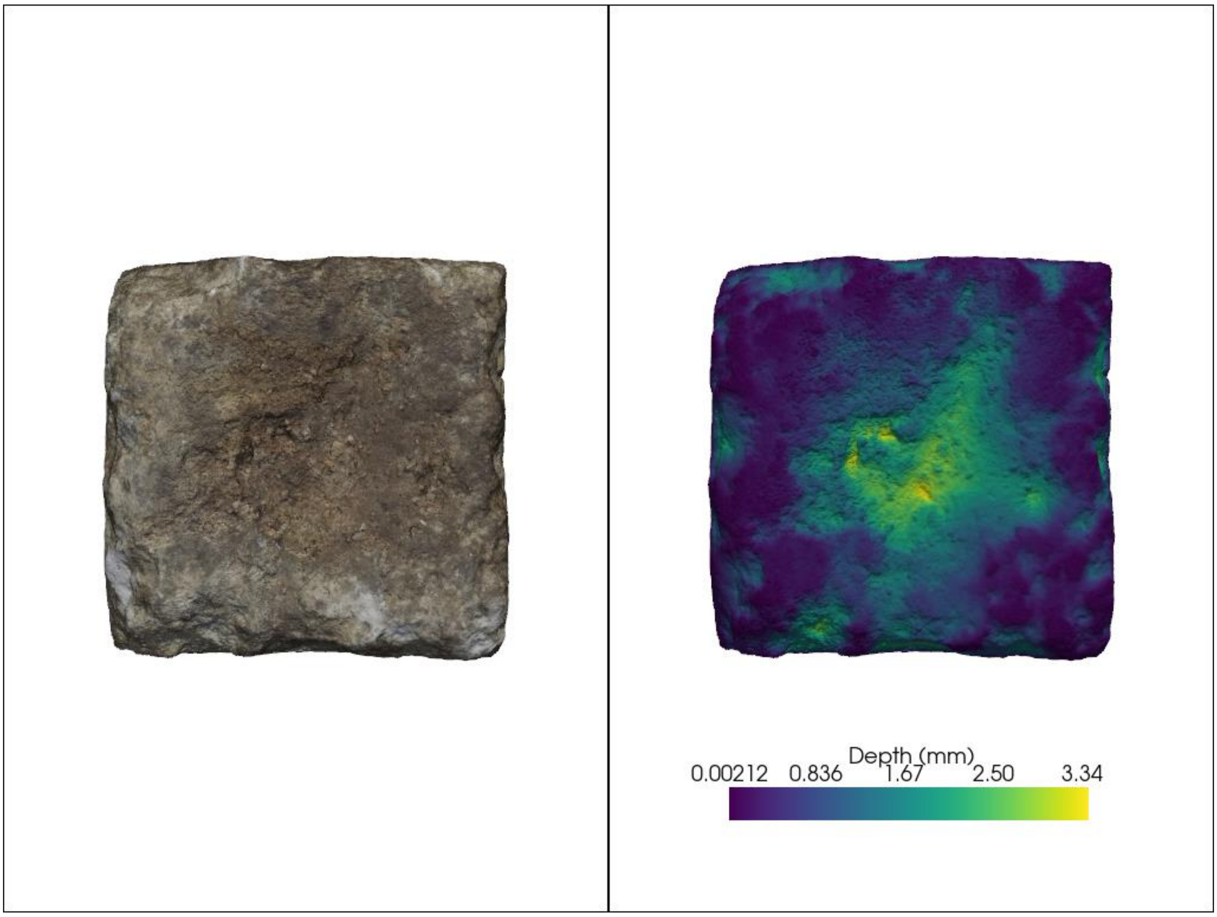


Fig S1: A pitted limestone tool as shown in figure 1 of the main text exhibiting a pit (Left) and the quantification of depth using the distance to convex hull method outlined in Proffitt et. al. (2021)."
