## Supplementary Model Information for "Social cues on stone tools outweigh raw material properties in wild primates"

### 1 Task 1

bystanders: 344 trials (1261 trials no/NA)

#### 1.1 Anvil selection

```
> # full-reduced model comparison: effect bystanders
> as.data.frame(anova(LM.selection.A.full.wnc.by, LM.selection.A.full.wnc,
test="Chisq"))
                           npar      AIC      BIC    logLik deviance    Chi
sq Df Pr(>Chisq)
LM.selection.A.full.wnc      37 777.6337 940.8102 -351.8168 703.6337
NA NA         NA
LM.selection.A.full.wnc.by   38 778.1245 945.7112 -351.0623 702.1245 1.5091
43  1    0.21927


> # results bystanders
> LM.selection.A.full.wnc.by.sum <- summary(LM.selection.A.full.wnc.by)
> round(LM.selection.A.full.wnc.by.sum$coefficients, 3)
                 Estimate Std. Error z value Pr(>|z|)
(Intercept)        -0.015      0.352  -0.044    0.965
materialsandstone  -0.687      0.273  -2.512    0.012
positionL2          1.188      0.410   2.900    0.004
positionR1         -0.575      0.492  -1.168    0.243
positionR2          0.474      0.372   1.272    0.203
sexM               -0.218      0.260  -0.836    0.403
parkSCNP           -0.886      0.296  -2.990    0.003
z.rel.trial        -0.124      0.137  -0.905    0.365
bystandersyes       0.313      0.251   1.246    0.213
> LM.selection.A.full.wnc.by.drop1 <- as.data.frame(round(drop1(LM.selectio
n.A.full.wnc.by, test="Chisq"), 3))
boundary (singular) fit: see help('isSingular')
boundary (singular) fit: see help('isSingular')
boundary (singular) fit: see help('isSingular')
boundary (singular) fit: see help('isSingular')
boundary (singular) fit: see help('isSingular')
boundary (singular) fit: see help('isSingular')
> LM.selection.A.full.wnc.by.drop1
            npar     AIC   LRT Pr(Chi)
<none>        NA 778.125    NA      NA
material       1 782.537 6.413   0.011
position       3 780.306 8.182   0.042
sex            1 776.811 0.687   0.407
park           1 783.925 7.801   0.005
z.rel.trial    1 776.893 0.768   0.381
bystanders     1 777.634 1.509   0.219
```

#### 1.2 Hammer selection

```
> # full-reduced model comparison: effect bystanders
> as.data.frame(anova(LM.selection.H.full.wnc.by, LM.selection.H.full.wnc,
test="Chisq"))
                           npar      AIC      BIC    logLik deviance    Chi
sq Df Pr(>Chisq)
LM.selection.H.full.wnc      37 735.7991 898.9755 -330.8995 661.7991
NA NA         NA
```

```
LM.selection.H.full.wnc.by   38 736.5418 904.1285 -330.2709 660.5418 1.2572
27  1  0.2621766


> # results bystanders
> LM.selection.H.full.wnc.by.sum <- summary(LM.selection.H.full.wnc.by)
> round(LM.selection.H.full.wnc.by.sum$coefficients, 3)
                  Estimate Std. Error z value Pr(>|z|)
(Intercept)         -1.214      0.416  -2.917    0.004
materialsandstone    0.783      0.348   2.249    0.025
positionL2          -0.385      0.456  -0.846    0.398
positionR1          -1.282      0.471  -2.719    0.007
positionR2           0.157      0.407   0.386    0.699
sexM                 0.174      0.268   0.646    0.518
parkSCNP            -0.290      0.325  -0.891    0.373
z.rel.trial          0.062      0.122   0.504    0.614
bystandersyes        0.124      0.274   0.454    0.650
> LM.selection.H.full.wnc.by.drop1 <- as.data.frame(round(drop1(LM.selectio
n.H.full.wnc.by, test="Chisq"), 3))
boundary (singular) fit: see help('isSingular')
boundary (singular) fit: see help('isSingular')
boundary (singular) fit: see help('isSingular')
boundary (singular) fit: see help('isSingular')
boundary (singular) fit: see help('isSingular')
boundary (singular) fit: see help('isSingular')
> LM.selection.H.full.wnc.by.drop1
            npar     AIC    LRT Pr(Chi)
<none>        NA 736.542     NA      NA
material       1 740.757  6.215   0.013
position       3 746.400 15.859   0.001
sex            1 736.087  1.546   0.214
park           1 735.591  1.049   0.306
z.rel.trial    1 734.799  0.257   0.612
bystanders     1 735.799  1.257   0.262
```

### 2 Task 2

site: bystanders: 154 trials (288 trials no/NA)

anvil/hammer: bystanders: 629 trials (1195 trials no/NA)

#### 2.1 Site selection

```
> as.data.frame(anova(LM.selection.first.full.wnc.by, LM.selection.first.fu
ll.wnc, test="Chisq"))
                              npar      AIC      BIC    logLik deviance Ch
isq Df Pr(>Chisq)
LM.selection.first.full.wnc      17 516.3048 585.8571 -241.1524 482.3048
NA  NA          NA
LM.selection.first.full.wnc.by   18 518.3048 591.9484 -241.1524 482.3048
0  1           1


> # results bystanders
> LM.selection.first.full.wnc.by.sum <- summary(LM.selection.first.full.wnc
.by)
> round(LM.selection.first.full.wnc.by.sum$coefficients, 3)
               Estimate Std. Error z value Pr(>|z|)
```

```
(Intercept)         0.930      0.401   2.318    0.020
siteraw_stones     -1.820      0.456  -3.991    0.000
sideright          -0.039      0.702  -0.056    0.955
sexM                0.000      0.255   0.000    1.000
parkSCNP            0.000      0.248   0.000    1.000
rel.trial           0.000      0.027   0.000    1.000
bystandersyes       0.000      0.267   0.000    1.000
> LM.selection.first.full.wnc.by.drop1 <- as.data.frame(round(drop1(LM.sele
ction.first.full.wnc.by, test="Chisq"), 3))
boundary (singular) fit: see help('isSingular')
boundary (singular) fit: see help('isSingular')
boundary (singular) fit: see help('isSingular')
boundary (singular) fit: see help('isSingular')
boundary (singular) fit: see help('isSingular')
boundary (singular) fit: see help('isSingular')
> LM.selection.first.full.wnc.by.drop1
           npar     AIC   LRT Pr(Chi)
<none>       NA 518.305    NA      NA
site          1 525.742 9.437   0.002
side          1 516.307 0.002   0.961
sex           1 516.305 0.000   1.000
park          1 516.305 0.000   1.000
rel.trial     1 516.305 0.000   1.000
bystanders    1 516.305 0.000   1.000
```

#### 2.2 Anvil selection

```
> as.data.frame(anova(LM.selection.A.full.wnc.by, LM.selection.A.full.wnc,
test="Chisq"))
                           npar      AIC      BIC    logLik deviance     Ch
isq Df Pr(>Chisq)
LM.selection.A.full.wnc      46 948.4119 1161.302 -428.2059 856.4119
NA  NA         NA
LM.selection.A.full.wnc.by   47 950.2013 1167.719 -428.1007 856.2013 0.2105
676   1  0.6463229


> # results bystanders
> LM.selection.A.full.wnc.by.sum <- summary(LM.selection.A.full.wnc.by)
> round(LM.selection.A.full.wnc.by.sum$coefficients, 3)
                  Estimate Std. Error z value Pr(>|z|)
(Intercept)         -0.141      0.393  -0.359    0.720
materialsandstone   -0.990      0.246  -4.028    0.000
percussivetool       1.334      0.351   3.798    0.000
positionL2           0.189      0.364   0.519    0.604
positionR1           0.026      0.445   0.058    0.954
positionR2           0.484      0.523   0.924    0.356
sexM                 0.120      0.256   0.471    0.638
parkSCNP            -1.333      0.284  -4.698    0.000
z.rel.trial         -0.045      0.120  -0.378    0.705
bystandersyes       -0.104      0.227  -0.459    0.646
> LM.selection.A.full.wnc.by.drop1 <- as.data.frame(round(drop1(LM.selectio
n.A.full.wnc.by, test="Chisq"), 3))
boundary (singular) fit: see help('isSingular')
boundary (singular) fit: see help('isSingular')
boundary (singular) fit: see help('isSingular')
boundary (singular) fit: see help('isSingular')
boundary (singular) fit: see help('isSingular')
boundary (singular) fit: see help('isSingular')
boundary (singular) fit: see help('isSingular')
> LM.selection.A.full.wnc.by.drop1
            npar     AIC    LRT Pr(Chi)
<none>        NA 950.201     NA      NA
material       1 964.082 15.880   0.000
percussive     1 960.154 11.953   0.001
position       3 945.259  1.058   0.787
sex            1 948.421  0.220   0.639
```

```
park            1 970.211 22.009    0.000
z.rel.trial     1 950.926  2.724    0.099
bystanders      1 948.412  0.211    0.646
```

#### 2.3 Hammer selection

```
> # full-reduced model comparison: effect bystanders
> as.data.frame(anova(LM.selection.H.full.wnc.by, LM.selection.H.full.wnc,
test="Chisq"))
                           npar      AIC      BIC    logLik deviance Chisq
Df Pr(>Chisq)
LM.selection.H.full.wnc      46 855.4767 1068.367 -381.7384 763.4767    NA
NA         NA
LM.selection.H.full.wnc.by   47 857.5924 1075.110 -381.7962 763.5924     0
1          1


> # results bystanders
> LM.selection.H.full.wnc.by.sum <- summary(LM.selection.H.full.wnc.by)
> round(LM.selection.H.full.wnc.by.sum$coefficients, 3)
                  Estimate Std. Error z value Pr(>|z|)
(Intercept)         -2.240      0.510  -4.389    0.000
materialsandstone    0.745      0.350   2.125    0.034
percussivetool       1.392      0.446   3.123    0.002
positionL2          -0.444      0.563  -0.788    0.431
positionR1           0.419      0.506   0.828    0.408
positionR2          -0.841      0.558  -1.506    0.132
sexM                 0.174      0.272   0.641    0.522
parkSCNP            -0.358      0.305  -1.172    0.241
z.rel.trial          0.012      0.132   0.087    0.930
bystandersyes        0.020      0.270   0.075    0.940
> LM.selection.H.full.wnc.by.drop1 <- as.data.frame(round(drop1(LM.selectio
n.H.full.wnc.by, test="Chisq"), 3))
boundary (singular) fit: see help('isSingular')
boundary (singular) fit: see help('isSingular')
boundary (singular) fit: see help('isSingular')
boundary (singular) fit: see help('isSingular')
boundary (singular) fit: see help('isSingular')
boundary (singular) fit: see help('isSingular')
boundary (singular) fit: see help('isSingular')
> LM.selection.H.full.wnc.by.drop1
            npar     AIC    LRT Pr(Chi)
<none>        NA 857.592     NA      NA
material       1 859.887  4.294   0.038
percussive     1 865.284  9.692   0.002
position       3 854.428  2.836   0.418
sex            1 855.910  0.318   0.573
park           1 856.767  1.175   0.278
z.rel.trial    1 855.481 -0.112   1.000
bystanders     1 855.477 -0.116   1.000
```

### 3 Task 3

anvil/hammer: bystanders: 368 trials (234 trials no/NA)

#### 3.1 Anvil selection

```
> # full-reduced model comparison: effect bystanders
```

```
> as.data.frame(anova(LM.selection.A.full.wnc.by, LM.selection.A.full.wnc,
test="Chisq"))
                           npar      AIC      BIC    logLik deviance      C
hisq Df Pr(>Chisq)
LM.selection.A.full.wnc      46 403.4970 572.6277 -155.7485 311.4970
NA NA          NA
LM.selection.A.full.wnc.by   47 405.4743 578.2817 -155.7371 311.4743 0.0227
3152  1  0.8801574


> # results bystanders
> LM.selection.A.full.wnc.by.sum <- summary(LM.selection.A.full.wnc.by)
> round(LM.selection.A.full.wnc.by.sum$coefficients, 3)
                  Estimate Std. Error z value Pr(>|z|)
(Intercept)          0.261      1.108   0.236    0.813
materialsandstone   -0.638      0.533  -1.196    0.232
percussivetool       1.527      0.459   3.327    0.001
positionL2          -0.926      1.082  -0.856    0.392
positionR1          -2.022      0.983  -2.056    0.040
positionR2          -1.406      0.942  -1.493    0.136
sexM                 0.717      0.499   1.436    0.151
parkSCNP            -0.380      0.889  -0.427    0.669
z.rel.trial          0.210      0.199   1.053    0.292
bystandersyes       -0.058      0.386  -0.151    0.880
> LM.selection.A.full.wnc.by.drop1 <- as.data.frame(round(drop1(LM.selectio
n.A.full.wnc.by, test="Chisq"), 3))
boundary (singular) fit: see help('isSingular')
boundary (singular) fit: see help('isSingular')
boundary (singular) fit: see help('isSingular')
boundary (singular) fit: see help('isSingular')
boundary (singular) fit: see help('isSingular')
boundary (singular) fit: see help('isSingular')
boundary (singular) fit: see help('isSingular')
> LM.selection.A.full.wnc.by.drop1
            npar     AIC    LRT Pr(Chi)
<none>        NA 405.474     NA      NA
material       1 404.940  1.465   0.226
percussive     1 414.212 10.737   0.001
position       3 405.504  6.030   0.110
sex            1 405.714  2.240   0.134
park           1 403.655  0.180   0.671
z.rel.trial    1 404.686  1.211   0.271
bystanders     1 403.497  0.023   0.880
```

#### 3.2 Hammer selection

```
> # full-reduced model comparison: effect bystanders
> as.data.frame(anova(LM.selection.H.full.wnc.by, LM.selection.H.full.wnc,
test="Chisq"))
                           npar      AIC      BIC    logLik deviance     Ch
isq Df Pr(>Chisq)
LM.selection.H.full.wnc      46 409.1068 578.2375 -158.5534 317.1068
NA NA          NA
LM.selection.H.full.wnc.by   47 410.1280 582.9354 -158.0640 316.1280 0.9788
263  1  0.3224887

> # results bystanders
> LM.selection.H.full.wnc.by.sum <- summary(LM.selection.H.full.wnc.by)
> round(LM.selection.H.full.wnc.by.sum$coefficients, 3)
                  Estimate Std. Error z value Pr(>|z|)
(Intercept)         -3.197      1.463  -2.186    0.029
materialsandstone    0.704      0.591   1.191    0.234
percussivetool      -0.613      0.449  -1.367    0.172
positionL2           2.124      1.266   1.677    0.094
positionR1           0.855      0.918   0.932    0.351
positionR2           1.473      1.081   1.362    0.173
sexM                 0.195      0.477   0.410    0.682
```

```
parkSCNP               1.011      0.951   1.063    0.288
z.rel.trial           -0.086      0.215  -0.400    0.689
bystandersyes         -0.405      0.410  -0.987    0.323
> LM.selection.H.full.wnc.by.drop1 <- as.data.frame(round(drop1(LM.selectio
n.H.full.wnc.by, test="Chisq"), 3))
boundary (singular) fit: see help('isSingular')
boundary (singular) fit: see help('isSingular')
boundary (singular) fit: see help('isSingular')
boundary (singular) fit: see help('isSingular')
boundary (singular) fit: see help('isSingular')
boundary (singular) fit: see help('isSingular')
boundary (singular) fit: see help('isSingular')
Warnmeldungen:
1: In optwrap(optimizer, devfun, start, rho$lower, control = control,  :
  convergence code 1 from bobyqa: bobyqa -- maximum number of function eval
uations exceeded
2: In optwrap(optimizer, devfun, start, rho$lower, control = control,  :
  convergence code 1 from bobyqa: bobyqa -- maximum number of function eval
uations exceeded
> LM.selection.H.full.wnc.by.drop1
            npar     AIC   LRT Pr(Chi)
<none>        NA 410.128    NA      NA
material       1 409.719 1.591   0.207
percussive     1 409.984 1.856   0.173
position       3 407.075 2.947   0.400
sex            1 408.297 0.169   0.681
park           1 409.492 1.364   0.243
z.rel.trial    1 408.299 0.171   0.679
bystanders     1 409.107 0.979   0.322
```

### 4 Performance

Psuccess: bystanders: 146 trials (259 trials no/NA)

Lsuccess/Nstrikes: bystanders: 118 trials (174 trials no/NA)

#### 4.1 Success probability

```
> as.data.frame(anova(LM.Psuccess.full.wnc.by, LM.Psuccess.full.wnc, test="
Chisq"))
                        npar      AIC      BIC    logLik deviance    Chisq D
f Pr(>Chisq)
LM.Psuccess.full.wnc      25 473.5344 573.6315 -211.7672 423.5344       NA N
A         NA
LM.Psuccess.full.wnc.by   26 475.3087 579.4098 -211.6544 423.3087 0.22564
1  0.6347758

> # results bystanders
> LM.Psuccess.full.wnc.by.sum <- summary(LM.Psuccess.full.wnc.by)
> round(LM.Psuccess.full.wnc.by.sum$coefficients, 3)
                     Estimate Std. Error z value Pr(>|z|)
(Intercept)             0.027      0.489   0.056    0.955
A.materialsandstone     0.001      0.381   0.003    0.998
H.materialsandstone     0.141      0.345   0.408    0.683
z.abs.trial             0.498      0.179   2.785    0.005
sexM                   -0.123      0.456  -0.269    0.788
parkSCNP                1.530      0.448   3.419    0.001
bystandersyes           0.156      0.328   0.475    0.635
> LM.Psuccess.full.wnc.by.drop1 <- as.data.frame(round(drop1(LM.Psuccess.fu
ll.wnc.by, test="Chisq"), 3))
boundary (singular) fit: see help('isSingular')
```

```
boundary (singular) fit: see help('isSingular')
boundary (singular) fit: see help('isSingular')
boundary (singular) fit: see help('isSingular')
boundary (singular) fit: see help('isSingular')
boundary (singular) fit: see help('isSingular')
> LM.Psuccess.full.wnc.by.drop1
            npar     AIC    LRT Pr(Chi)
<none>        NA 475.309     NA      NA
A.material     1 473.309  0.000   0.998
H.material     1 473.475  0.166   0.683
z.abs.trial    1 480.646  7.338   0.007
sex            1 473.381  0.072   0.789
park           1 486.399 13.091   0.000
bystanders     1 473.534  0.226   0.635
```

#### 4.2 Success latency

```
> as.data.frame(anova(LM.Lsuccess.full.wnc.by, LM.Lsuccess.full.wnc, test="
Chisq"))
refitting model(s) with ML (instead of REML)
                        npar      AIC      BIC    logLik deviance       Chi
sq Df Pr(>Chisq)
LM.Lsuccess.full.wnc      25 817.1960 909.1148 -383.5980 767.1960
NA  NA         NA
LM.Lsuccess.full.wnc.by   26 819.1918 914.7874 -383.5959 767.1918 0.0041594
82  1  0.9485768

> # results bystanders
> LM.Lsuccess.full.wnc.by.sum <- summary(LM.Lsuccess.full.wnc.by)
> round(LM.Lsuccess.full.wnc.by.sum$coefficients, 3)
                     Estimate Std. Error t value
(Intercept)             2.614      0.258  10.137
A.materialsandstone     0.206      0.150   1.373
H.materialsandstone     0.007      0.125   0.054
z.abs.trial            -0.030      0.078  -0.382
sexM                   -0.010      0.240  -0.040
parkSCNP               -0.095      0.253  -0.376
bystandersyes          -0.008      0.120  -0.069
> LM.Lsuccess.full.wnc.by.drop1 <- as.data.frame(round(drop1(LM.Lsuccess.fu
ll.wnc.by, test="Chisq"), 3))
boundary (singular) fit: see help('isSingular')
boundary (singular) fit: see help('isSingular')
boundary (singular) fit: see help('isSingular')
boundary (singular) fit: see help('isSingular')
boundary (singular) fit: see help('isSingular')
boundary (singular) fit: see help('isSingular')
> LM.Lsuccess.full.wnc.by.drop1
            npar     AIC   LRT Pr(Chi)
<none>        NA 819.192    NA      NA
A.material     1 818.913 1.721   0.190
H.material     1 817.194 0.002   0.964
z.abs.trial    1 817.368 0.176   0.675
sex            1 817.192 0.000   0.988
park           1 817.293 0.102   0.750
bystanders     1 817.196 0.004   0.949
```

#### 4.3 Number of strikes

```
> # full-reduced model comparison: effect bystanders
> as.data.frame(anova(LM.Nstrikes.full.wnc.by, LM.Nstrikes.full.wnc, test="
Chisq"))
```

```
                            npar      AIC      BIC    logLik deviance      Chis
q Df Pr(>Chisq)
LM.Nstrikes.full.wnc      25 1548.481 1640.400 -749.2406 1498.481         N
A NA        NA
LM.Nstrikes.full.wnc.by   26 1550.436 1646.031 -749.2179 1498.436 0.0454782
5  1   0.831127

> # results bystanders
> LM.Nstrikes.full.wnc.by.sum <- summary(LM.Nstrikes.full.wnc.by)
> round(LM.Nstrikes.full.wnc.by.sum$coefficients, 3)
                    Estimate Std. Error z value Pr(>|z|)
(Intercept)            1.255      0.198   6.342    0.000
A.materialsandstone    0.078      0.120   0.650    0.516
H.materialsandstone   -0.005      0.118  -0.045    0.964
z.abs.trial            0.032      0.063   0.507    0.612
sexM                   0.081      0.183   0.445    0.656
parkSCNP               0.309      0.197   1.570    0.116
bystandersyes         -0.019      0.087  -0.216    0.829
> LM.Nstrikes.full.wnc.by.drop1 <- as.data.frame(round(drop1(LM.Nstrikes.fu
ll.wnc.by, test="Chisq"), 3))
boundary (singular) fit: see help('isSingular')
boundary (singular) fit: see help('isSingular')
boundary (singular) fit: see help('isSingular')
boundary (singular) fit: see help('isSingular')
boundary (singular) fit: see help('isSingular')
boundary (singular) fit: see help('isSingular')
> LM.Nstrikes.full.wnc.by.drop1
            npar      AIC   LRT Pr(Chi)
<none>        NA 1550.436    NA      NA
A.material     1 1548.855 0.419   0.517
H.material     1 1548.438 0.002   0.965
z.abs.trial    1 1548.700 0.264   0.607
sex            1 1548.625 0.189   0.664
park           1 1550.827 2.391   0.122
bystanders     1 1548.481 0.045   0.831
```
